# Adipose gene expression profiles reveal novel insights into the adaptation of northern Eurasian semi-domestic reindeer (*Rangifer tarandus*)

**DOI:** 10.1101/2021.04.17.440269

**Authors:** Melak Weldenegodguad, Kisun Pokharel, Laura Niiranen, Päivi Soppela, Innokentyi Ammosov, Mervi Honkatukia, Heli Lindeberg, Jaana Peippo, Tiina Reilas, Nuccio Mazzullo, Kari A. Mäkelä, Tommi Nyman, Arja Tervahauta, Karl-Heinz Herzig, Florian Stammler, Juha Kantanen

## Abstract

Reindeer (*Rangifer tarandus*) are semi-domesticated animals adapted to the challenging arctic conditions of northern Eurasia. Adipose tissues play a crucial role in animals living in northern environments by altering gene expression in their tissues to regulate energy homeostasis and thermogenic activity. Here, we performed transcriptome profiling by RNA sequencing of adipose tissues from three different anatomical depots: metacarpal (bone marrow), perirenal, and prescapular fat in Finnish and Even reindeer (in Sakha) during two seasonal time points (spring and winter). On average 36.5 million pair-ended clean reads were obtained for each sample, and a total of 16,362 genes were expressed in our data. Gene expression profiles in metacarpal tissue were distinct and clustered separately from perirenal and prescapular adipose tissues. Notably, metacarpal adipose tissue appeared to have a significant role in the regulation of the energy metabolism of reindeer in spring when their nutritional condition is poor after winter. During spring, when the animals are in less optimal condition, genes associated with the immune system (e.g., *CCL2*, *CCL11*, *CXCL14*, *IGSF3*, *IGHM*, *IGLC7*, *IGKC*, *JCHAIN,* and *IGSF10*) were upregulated in the perirenal and prescapular adipose tissue, while genes involved in energy metabolism (e.g., *ACOT2*, *APOA1*, *ANGPTL1*, *ANGPTL8*, *ELOVL7*, *MSMO1*, *PFKFB1*, and *ST3GAL6*) were upregulated in metacarpal tissue. Even reindeer harboured relatively fewer significantly differentially expressed genes than Finnish reindeer, irrespective of the season, possibly owing to climatic and management differences. Moreover, blood and tissue parameters reflecting general physiological and metabolic status showed less seasonal variation in Even reindeer than in Finnish reindeer. This study identified adipose candidate genes potentially involved in immune response, fat deposition, energy metabolism, development, cell growth, and organogenesis. Taken together, this study provides new information on the mechanisms by which reindeer adapt to less optimal arctic conditions.

## Introduction

Native to northern and subarctic regions of Eurasia, reindeer (*Rangifer tarandus*) have societal, cultural, and ecological values for the livelihoods of arctic indigenous people and pastoralists, and have multiple socio-economic roles, such as providing meat, hides, milk, and serving as a means of transportation [1–3]. Reindeer survive in challenging northern and extreme arctic environments characterized by low temperatures, prolonged daylight during summers, and darkness and limited availability of grazing resources during long winters [3, 4]. Adipose tissues are vital for reindeer to adapt to such extreme conditions [5, 6]. Adipose tissues are important organs for several functions in energy metabolism that are crucial for survival and successful reproduction. White adipose tissue (WAT) stores energy in the form of lipids and serves as a long-term energy reserve, whereas brown adipose tissue (BAT) contributes to both thermohomeostasis and energy balance by producing heat via the function of uncoupling protein 1 (*UCP1*) [7]. WAT is also an important endocrine organ that secretes several hormones, including adipomyokines and cytokines, which contribute to energy metabolism and immunity and act as signals to the central nervous system [8, 9]. The reindeer is a lean animal, but it also relies on WAT as a source of energy and hormonal signals in changing environmental conditions [5]. In newborn reindeer, BAT plays a crucial role in regulating non-shivering thermogenesis, but its effect diminishes over time [10, 11]. However, browning of WAT has been observed in various species during later stages of life after chronic cold exposure [12] and due to pharmacological and nutritional agents [13]. Bone marrow has a specific type of adipose tissue (BMAT) that acts as an energy reservoir, contributes to local and systemic metabolic processes [14, 15], and undergoes dynamic changes [16]. BMAT decreases as a result of starvation in freely grazing animals [17–19].

The majority of adipose transcriptome studies using high-throughput RNA sequencing (RNA-Seq) have been limited to mice [20, 21], sheep [22–24], pigs [25–27], and humans [28–30]. Adipose transcriptomes are affected by the type of adipose, as well as by the sex and age of the animal [31]. Changes in gene expression within adipose tissues in response to temperature fluctuations have been studied mostly in mice so far [20,21,32]. However, gene expression profiles in reindeer adipose tissues have thus far not been investigated. Adipose tissues from various parts of the body have their own unique functions. In the present study, we investigated gene expression profiles of three adipose tissue depots: metacarpal (M), perirenal (P), and prescapular (S) tissues. These adipose depots were selected because they represent visceral (P), peripheral (S), and bone marrow (M) fat. These anatomical depots are also expected to reflect different metabolic functions. For example, the prescapular area is a major BAT depot in newborn reindeer [11] and thus an interesting target for regulating the expression of *UCP1*, as well as other markers for cold adaptation. The samples were collected from the Finnish and Even (Sakha Republic, Yakutia, Russia) reindeer breeds (or populations), which belong to two different phylogenetic clusters of Eurasian reindeer (Pokharel K. et al., unpublished) and differ in present-day management and feeding practices. In Yakutia, reindeer feed on natural pastures with expressed seasonal variation and high migratory behaviour (high lichen content in winter, fresh leaves in spring, grass, and mountain herbs in summer), whereas in Finland, extra fodder (concentrates) is provided during peak winter (February and March). Reindeer herders observe that the lichen-rich diet in winter helps reindeer keep their weight and survive the cold, while the grassy diet in summer accounts for the principal annual gain in weight. In Yakutia, reindeer have to survive in extreme climatic conditions where the annual temperature fluctuates from -60°C up to more than +30°C.

Here, we aim to obtain insights into the seasonal fluctuations in the transcriptome profiles of three adipose tissues (metacarpal, perirenal, and prescapular) in two reindeer populations (Even and Finnish). The tissue sampling was conducted during (early) winter (November–December) and (early) spring (April), when the animals were in the best and worst nutritional conditions, respectively. For the Even reindeer, we also compared the transcriptome profiles of these adipose tissues between male and female individuals. Moreover, to assess the health and physiological condition of the reindeer in different seasons and geographical locations, and to complement genetic analyses with phenotypic data, we analysed several blood metabolites along with hormones regulating energy metabolism, such as blood insulin, leptin, and hormone sensitive lipase (HSL), and proteins such as *UCP1* and *COX4* from adipose tissue. We expect that assessing changes in gene expression in reindeer adipose tissue due to seasonal variation may reveal adaptation mechanisms and help to understand the evolution of this adaptive response in reindeer and other mammalian species sharing northern Eurasian habitat conditions.

## Results

### RNA sequencing and mapping

A total of 220.5 gigabases (Gb) of RNA-seq data were generated from 56 adipose tissue samples collected from 19 Finnish and Even reindeer. As the adapters were trimmed automatically, and the Phred quality scores of the reads from all samples were greater than 30, we did not perform further trimming and quality filtering. The number of reads per sample ranged from 26.6 million (M) (4.0 Gb) to 217 M (32.6 Gb), with a mean of high-quality 36.5 M, 2 × 75 bp pair-ended reads per sample (S1 Table). As shown in S1 Table, the two samples (FR12_SCAP and YR1_SCAP), revealing the highest numbers of reads (217 M and 165 M, respectively), will be used to detect lncRNAs in a future study. The proportion of reads mapped to the reindeer reference genome ranged from 81% to 92%, with, on average, >90% of the reads from each sample uniquely mapped to the reindeer draft genome assembly (S2 Table). Raw sequence reads in compressed fastq format (fastq.gz) analyzed in this study have been deposited to the European Nucleotide Archive (ENA) and are publicly available under accession XXXXXX

### Gene expression overview

A total of 16,362 genes were expressed (cpm > 0.5 in at least two samples) in all 56 samples (S3 Table), representing approximately 60% of the 27,332 reindeer genes reported in the draft reindeer genome assembly annotation file [3]. The highest number of genes were expressed in metacarpal adipose tissue (n = 15,761), followed by prescapular (n = 15,087) and perirenal (n = 14,920) adipose tissues. Moreover, we examined the expressed genes across these three tissues to search for shared and uniquely expressed genes in the respective region and season (Fig 1). In all cohorts (spring and winter Finnish and Even samples), the highest number of expressed genes was found in the metacarpal adipose tissue. This tissue also displayed a higher number of tissue-specific expressed genes than did the perirenal and prescapular adipose tissues (Fig 1). In various region-season comparisons between the adipose tissues, >13,000 genes were commonly expressed. To assess expression similarity among the samples, we performed Principal Component Analysis PCA based on the top 500 most variable genes (Fig 2, S1 Fig). The PCA plot (Fig 2) shows that the metacarpal tissue samples were clustered separately from the other two tissues, and additional grouping of the samples along axis 2 based on two reindeer breeds. Similarly, a hierarchal clustering based on the top 25 genes with the highest variance across all samples also showed a similar pattern that clearly separated metacarpal tissue and the other tissues (S2 Fig).

**Fig 1.**
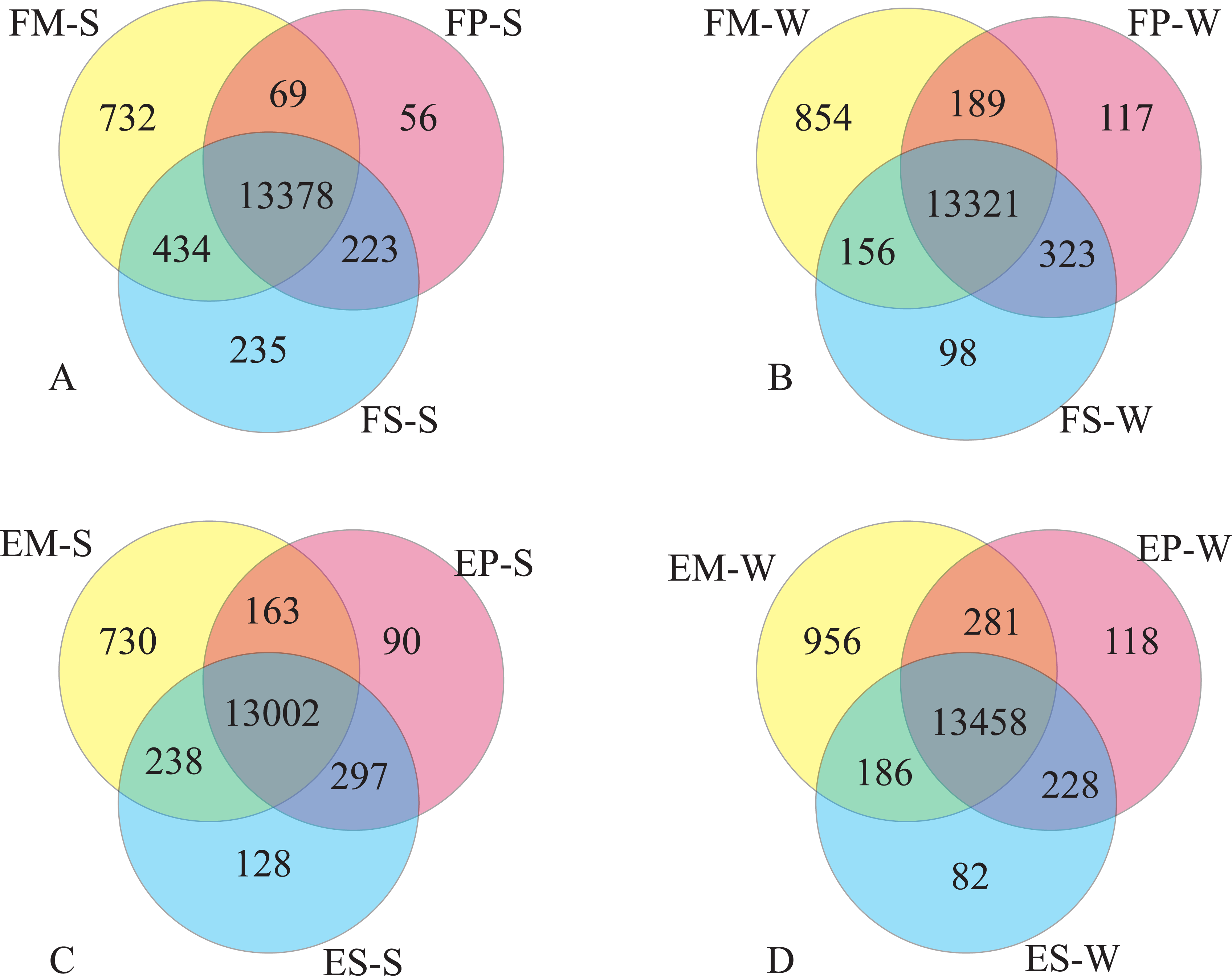
Venn diagram showing overlap of expressed genes (CPM ≥ 0.5 for at least two samples) among tissues in each region and season. Shared and uniquely expressed genes across tissues (M, P, and S) in (A) Finnish reindeer in the summer, (B) Finnish reindeer in the winter, (C) Even reindeer in the summer, and (D) Even reindeer in the winter.

**Fig 2.**
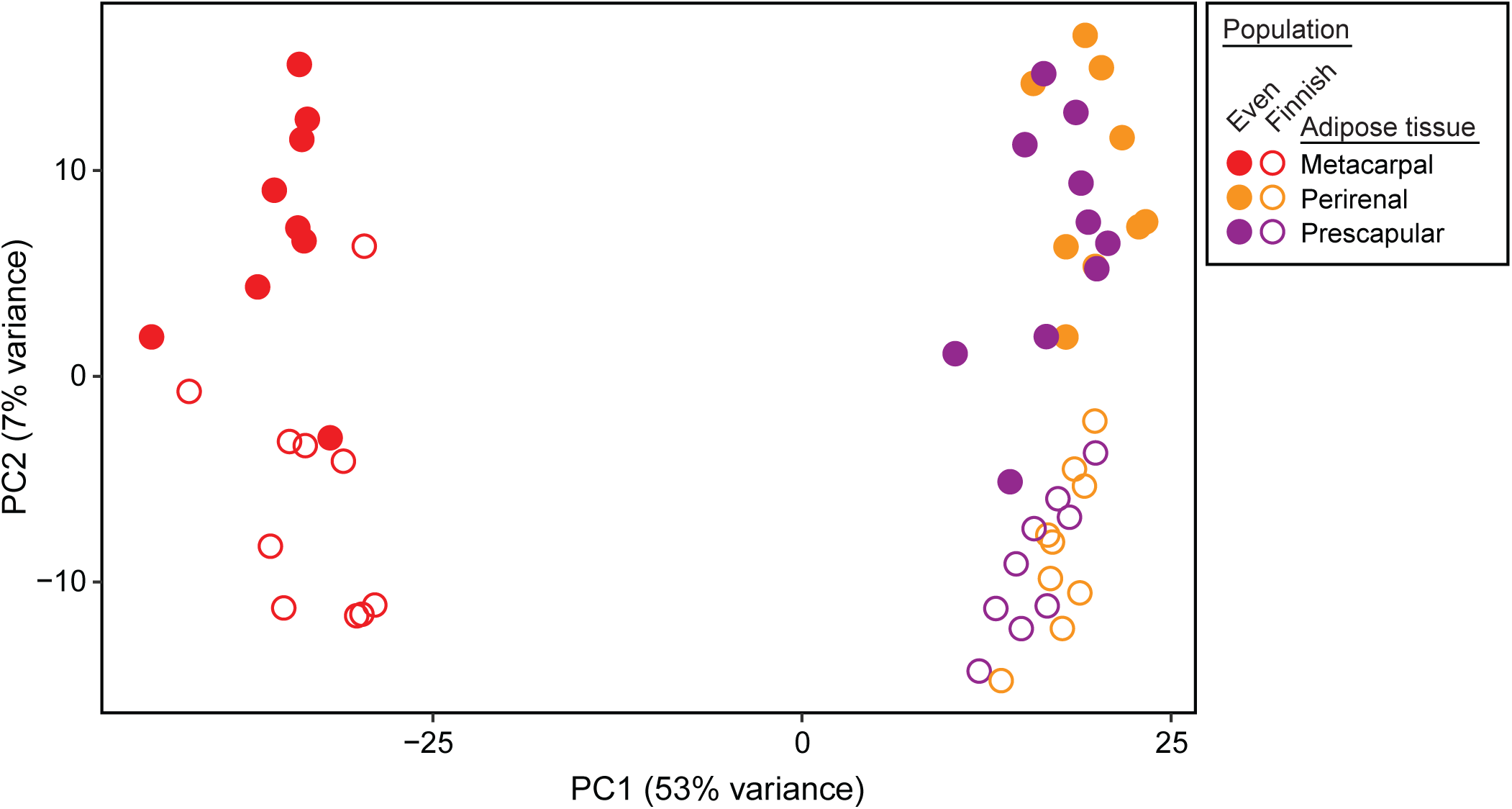
**PCA plots of the analyzed samples based on expression profiles, with dot colours indicating tissue and region (see legend).**

While more than 13,000 genes were commonly expressed in all tissues, several tissue-specific genes were identified in this study. In line with the distinct cluster observed in the PCA plot, more than 730 genes were uniquely expressed in the metacarpal adipose tissue (Fig 1). We further explored the genes specific to this tissue by removing the lowly expressed genes (TPM < 1). Out of 875 genes, 61 were commonly expressed in metacarpal adipose tissue in the different experimental groups. The metacarpal adipose tissue of Finnish reindeer collected in the winter had the largest number of uniquely expressed genes (n = 181), whereas Finnish metacarpal tissue collected in the spring possessed the lowest (n = 71) number of uniquely expressed genes. By contrast, Even reindeer samples collected in either season had roughly similar numbers of uniquely expressed genes (138 in spring and 136 in winter samples) (Fig 3).

**Fig 3.**
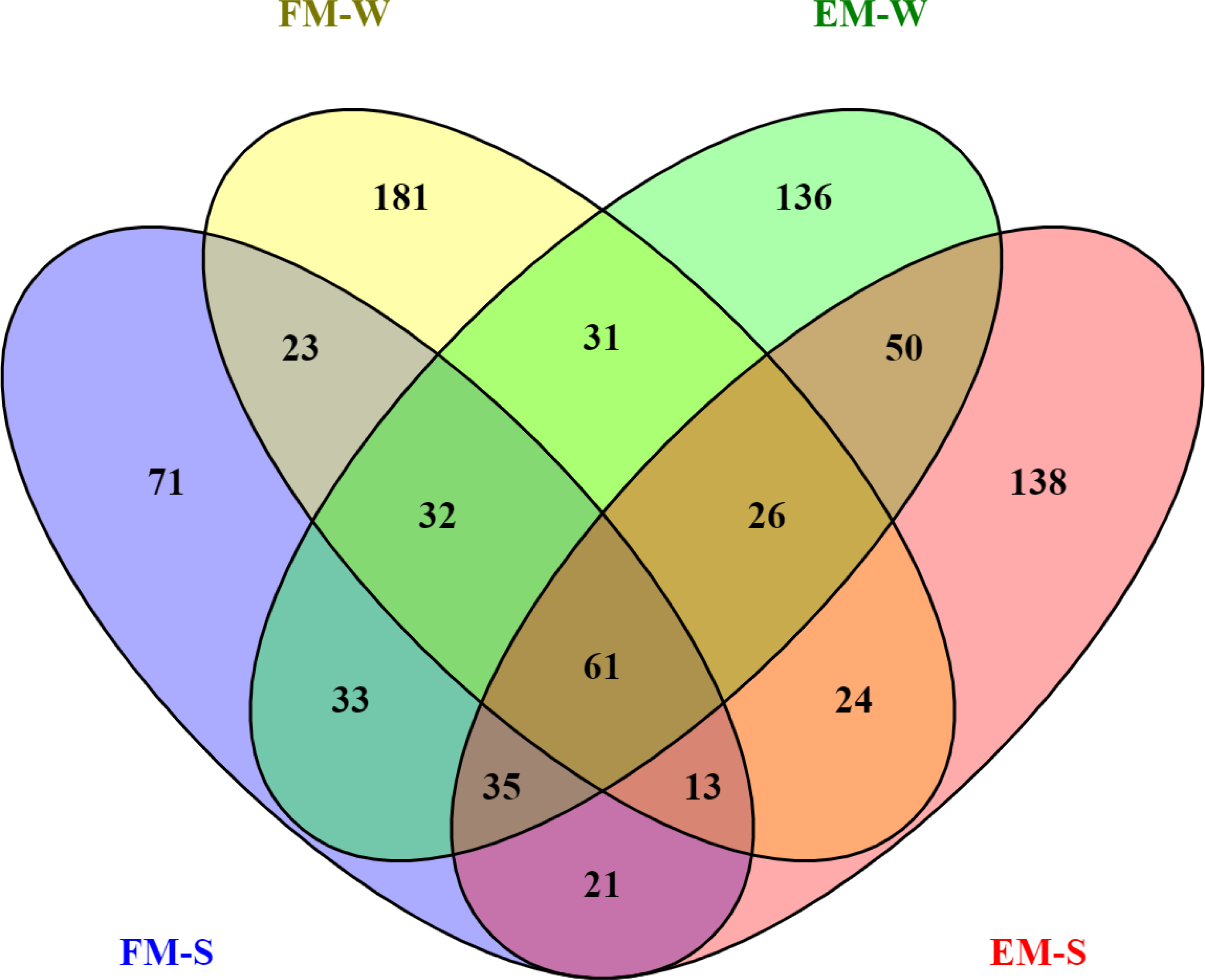
**Distribution of uniquely expressed genes in the metacarpal adipose tissue of Finnish (FM-W, FM-S) and Even (EM-W, EM-S) reindeer.**

Among the 61 genes that were unique to metacarpal adipose tissue yet shared by Finnish and Even reindeer, annotations were not available for 10 genes. The metacarpal-specific genes included several homeobox proteins (eg. *HOXD13*, *HOXA11*, *HOXD11*, *HOXA13*, *DLXC*), bone sialoprotein 2 (*IBSP*), osteomodulin (*OMD*), carbonic anhydrase 3 (*CA3*), C-X-C motif chemokine 10 (*CXCL10*), nuclear receptor-interacting protein 3 (*NRIP3*), and R-spondin-2 (*RSPO2)* (S4Table).

### Results from differential gene expression analyses

#### Seasonal differences in gene expression

For an insight into the effects of seasonal conditions, we compared gene expression profiles of adipose tissues from spring versus winter samples separately in the Finnish and Even reindeer. Finnish reindeer showed a higher number of significant differentially expressed genes (DEGs) than the Even reindeer (Table 2).

**Table 2.** The number of significantly identified differentially expressed genes (DEGs) in Finnish (F) and Even (E) reindeer due to seasonal changes (S, spring, W, winter) in in the metacarpal (M), perirenal (P), and prescapular (S) adipose tissues.

| Comparison | Total DEGs | Upregulated | Downregulated |
| --- | --- | --- | --- |
| FM-S vs. FM-W | 346 | 195 | 151 |
| FP-S vs. FP-W | 583 | 273 | 310 |
| FS-S vs. FS-W | 611 | 325 | 286 |
| EM-S vs. EM-W | 156 | 57 | 99 |
| EP-S vs. EP-W | 103 | 45 | 58 |
| ES-S vs. ES-W | 176 | 126 | 50 |

Altogether 346 genes were differentially expressed between seasons in metacarpal tissues of Finnish reindeer; of these, 195 were upregulated in spring samples and the rest were upregulated in the winter samples (Table 2, S5 Table and S4 Fig). Genes involved in metabolism (*ANGPTL8*, *BDH1*, *ASIP*, *QPRT*) and stress response (*BOLA3*, *GPR158*) were strongly upregulated in spring (LFC > 4) and those particularly associated with immune functions (*CCL11*, *CXCL8*, *CD33*, *IL1B*, *IRF4*) were strongly downregulated (LFC < -3.8) in spring metacarpal tissue (S5 Table and S4 Fig).

Among 583 DEGs in the perirenal tissue, genes such as *CCL11, TRH, MT1A, RPL38, AREG, NPW, SRSF3, RPS29, RPL36A* and *SLC11A1*, were highly upregulated (LFC > 4) in the spring samples and *MP68, RPL39, SLC39A12, SCN3A, RGS9, HSPA6, TNFRSF10B, EDIL3, RTP1* and *ABCC4* were highly downregulated (LFC < -3.7) (Table 2, S6 Table and S5 Fig). The DEGs upregulated in the spring samples include genes participating in the immune system (*SLC11A1*, *COMMD6*, *BATF*, *CCL19*), ribosomal and transcription processes (*RPL38*, *RPL36A*, *FKBP11*). On the other hand, downregulated genes were associated with signalling or signal transduction (*RGS9*, *RAPGEF5*, *COL4A5*, *PTGER2*, *MAML3*, *PDE5A*), cell differentiation or organogenesis (*GJA5*, *THSD7A*, *DACH1*, *MMRN2*, *NHSL2*, *ZFHX3*), and interaction of glucose and fatty acid metabolism (*PDK4*, SIK1).

The highest number (n = 611) of DEGs was found in prescapular tissue, 325 of which were upregulated in the spring samples (Table 2, S7 Table and S6 Fig). In prescapular adipose tissue from the spring, genes such as *SERPINA3-7, ESD, APOH, SERPINI2, SERPINC1, TCEB2, ITIH1, WNT9B, C8B, TRH, ASCL2, GSTA2* and *VPREB1* were among the top upregulated genes (LFC > 4.5) and *LHFP, TNFRSF10B, CRISP3, NEGR1, KCNA2, PTGER2, HSPA6, GTSF1* and *RPL30* were among the top downregulated genes (LFC < -3.5) (S7 Table and S6 Fig). A total of 30 genes, including *APOLD1, CYP26B1, IGFBP5, MICA, MICB, MT-ATP8, PDK4 and SCP2*, were commonly differentially expressed in three tissues (S3 Fig). Many of the genes in the prescapular tissue that were upregulated during spring were associated with inflammatory or immunological responses (*FXYD5*, *CCL19*, *IFIT3*, *CARD9*, *GMFG*, *SLC11A1*), feeding behaviour (*NPW*), adipogenesis (*DLK1*, *CITED4*), cell growth or differentiation (*MGP*, *ECSCR*, *TIMP1*, *MT1A*), and spermatogenesis (*SCHBP1L*, *MORN2*). Similarly, downregulated DEGs were associated with signalling (*RGS5*, *DCBLD2*, *RASGEF1B*, *RAPGEF5*), organogenesis (*PTGER2*, *KLF7*, *PHACTR2*, *KMT2A*, *DACH1*, *PDLIM5*), and lipid metabolism (*ARFGEF3*, *PDK4*, *FOXO1*, *PITPNC1*).

Similar pairwise comparison in Even reindeer revealed 156, 103, and 176 DEGs in metacarpal, perirenal, and prescapular tissues, respectively (Table 2, S8-S10 Tables, S8-S10 Figs). Prescapular adipose tissue had the highest number of unique significant DEGs (n = 141), followed by metacarpal (n = 135), and perirenal (n = 68) (S7 Fig). In both breeds, prescapular tissue harboured the highest (611 in Finnish and 176 in Even reindeer) number of DEGs (S7 and S10 Tables). Interestingly, while the number of upregulated genes was higher in the metacarpal tissue of Finnish samples from the winter collection, the same was not observed in Even reindeer. In the Even reindeer we, found seven common DEGs (*ORC5*, *ADI1, RPS15A, DAGLA, JCHAIN, ENSP00000353290*, and *FEZ1*) among all tissues (S7 Fig). DEGs that were upregulated in the spring samples of Even metacarpal adipose tissue appeared to have roles in lipid/fatty acid metabolism (*ELOVL7*, *APOA1*, *ACOT2*, *ANGPTL1*), cell structural functions (*CAPN6*, *MYL9*, *CLDN5*), and oxygen metabolism (*FMO1*, *FMO2*, *AOC1*, *STEAP1*). Downregulated DEGs in Even metacarpal adipose tissue were associated with development and organogenesis (*DMAP1*, *COL27A1*, *CA2*, *NRG3*, *JAK3*, *EPB41L3*, *ESM1*) and immune system (*C4A*, *IL17RB*, *HLA-DOA*, *SIGLEC1*, *CPA3*). In Even perirenal adipose tissue, several immunoglobulin related genes were upregulated during spring, suggesting activation of the immune system by pathogens. However, downregulated genes were mainly associated with lipid/energy metabolism (*ACACB*, *PRLR*, *APOL6*) and functions related to growth and development (*MEGF8*, *DAGLA*, *IGSF10*, *GRIP2*, *NSMF*). Among the three adipose tissues in Even reindeer, immune related DEGs (e.g., *JCHAIN*, *IGHM*, *IGKC*, *IGHV3-6*, *IGLC7*, *MUCM*) were predominantly upregulated in the prescapular tissue, whereas downregulated genes in the prescapular tissue were associated with lipid/energy metabolism (*TP53INP2*, *PPARGC1B*, *ACACB*) and growth and development (*DAGLA*, *IGS10*, *BMP5*, *FGFR2*, *NRG3*, *CYP26B1*).

#### Gene expression differences between the Finnish and Even reindeer

We made six pair-wise comparisons (three each for spring and winter samples) to identify DEGs between Finnish and Even reindeer (Table 3, S11-S16 Tables) of which the highest number (n = 504) of DEGs was present in metacarpal tissues collected during winter, whereas the same tissue revealed the lowest number (n = 126) of DEGs during spring (Table 2, S11 and S14 Tables). In comparisons involving spring samples, the highest numbers of DEGs were found in prescapular adipose tissue.

**Table 3.** The number of significantly identified DEGs in the metacarpal (M), perirenal (P), and prescapular (S) adipose tissues of Finnish (F) and Even (E) reindeer that were compared separately for spring (-S) and winter (-W) samples.

| Comparison | Total DEGs | Upregulated | Downregulated |
| --- | --- | --- | --- |
| EM-S vs FM-S | 126 | 78 | 48 |
| EP-S vs FP-S | 301 | 189 | 112 |
| ES-S vs FS-S | 385 | 229 | 156 |
| EM-W vs FM-W | 504 | 336 | 168 |
| EP-W vs FP-W | 401 | 182 | 219 |
| ES-W vs FS-W | 365 | 154 | 221 |

Four genes, *FIBP*, *CREB3L3*, *CLDN4,* and *ALKBH3,* were exclusively upregulated in the Finnish reindeer irrespective of the seasons. *FIBP* (*FGF1 Intracellular Binding Protein*) is known to promote mitogenic action to induce morphogenesis and differentiation. *CREB3L3* has been linked to triglyceride metabolism and growth suppression. *CLDN4* (Claudin 4) is a member of the claudin gene family. Being integral membrane proteins, claudins, in general, have a vital role in regulating the transport of solutes and ions through calcium-independent cell-adhesion activity.

Similarly, *TMEM182*, *AACS*, *FAM159B,* and *C19ORF80* were always upregulated in all five comparisons except between EM-S vs FM-S. Among the four genes, we did not find any relevant information about the function of *FAM159B*. There is relatively little information available on *TMEM182*, but its upregulation may be associated with adipose growth and remodelling [33, 34]. In the present context, a greater abundance of *TMEM182* in Even reindeer males might be linked to castration, as castrated males are known to accumulate more adipose tissues. *AACS* (*Acetoacetyl-CoA Synthetase*) appears to be involved in ketone body metabolism during adipose tissue development. *C19ORF80* (alternatively *ANGPTL8*, *Angiopoietin-like 8*) is known to mediate the transition between fasting and re-feeding and has an important role in the storage of fatty acids in adipose tissue during the refeeding state [35]. Interestingly, *C19ORF80* was downregulated in Even reindeer compared to Finnish reindeer among spring samples. We speculate that the additional winter feeding for the Finnish reindeer might have a role in the differential expression of *C19ORF80*.

#### Gender difference in gene expression

Our comparison of gene expression between female and male Even reindeer samples collected during winter revealed a total of 327 significant DEGs in the three adipose tissues (S17-S19 Tables).

We identified a total of 225 significant DEGs between female and male reindeer in metacarpal tissue (EM-F vs. EM-M) (Table 4 and S17 Table). Of 225 significant DEGs, 78 genes were upregulated, and 147 genes were downregulated in EM-F (Table 4 and S17 Table). The upregulated genes in female samples included *ANGPTL1, CX3CR1, CYP2B11, CYP2F3, CYP4B1, FABP6, ELOVL7, IGHV3-6, SLC19A3, SLC35F1* and *SLC6A17*, whereas genes such as *BMP1, BMP, IBSP, EFS, CCL19, IGDCC4, IL17RB, NAV3, NCAM1, NRL, OLFML2B, OLFML3, PRDM9, SCUBE1* and *FLT4* were downregulated.

**Table 4.** Summary of DEGS from male (-M) and female (-F) comparisons in metacarpal (M), perirenal (P), and prescapular (S) adipose tissues of Even (E) reindeer.

| Comparison | Total DEGs | Upregulated | Downregulated |
| --- | --- | --- | --- |
| EM-F vs EM-M | 225 | 78 | 147 |
| EP-F vs EP-M | 104 | 54 | 50 |
| ES-F vs ES-M | 49 | 27 | 22 |

In perirenal tissue, 104 significantly DEGs were detected between female and male reindeer, of which 54 and 50 genes were upregulated and downregulated in EP-F, respectively (Table 4 and S18 Table). The upregulated genes in female samples included *ABCG1, ACSL6, SLC14A1, SLC16A2, SLC4A10, SLC9A2, ZNF219, PLCD3, PTER, PLA2G5, ETNPPL, PLCD3, S1PR3,* and *S1PR3*, while *ATP5H, CCL3, COX7A1, CXCL9, NMB, NRL, PRDM9, SLC19A1, SLC22A5, ZFX,* and *ZRSR2* were downregulated.

Furthermore, 49 significantly DEGs were detected between female and male reindeer in prescapular tissue (ES-F vs. ES-M), of which 27 and 22 genes were upregulated and downregulated in ES-F, respectively (Table 4 and S19 Table). *GYS2, ARHGEF5, ABCC4, SUMO1, RPL39, SLC4A10*, and *ACSL6* were examples of upregulated genes in female samples, and genes such as *TXLNG, DDX3Y, USP9X, EIF2S3X, PRDM9, KDM6A, ZRSR2*, and *UCP1* were downregulated.

A total of 10 genes (*PRDM9, DDX3Y, ZRSR2, EIF2S3X, KDM6A, ZFX, UBA1, USP9X, TXLNG*, and *NRL*) were always upregulated in males irrespective of the adipose tissue type. These genes have diverse functions and may be linked to speciation (*PRDM9*), male infertility or spermatogenic failure (*DDX3Y*), mRNA splicing (*ZRSR2*), lipid metabolism (*EIF2S3X*), circadian rhythm (*USP9X*), cell cycle regulation (*TXLNG*), and photoreceptor development/function (*NRL*).

We did not find any genes commonly upregulated in all tissues of female samples; however, two upregulated DEGs (*CPE*, *RPL34*) were shared by metacarpal and perirenal adipose tissues, while 12 (*ARHGEF5*, *SLC4A10*, *ABCC4*, *SGCG*, *HAND1*, *CPXM2*, *GYS2*, *ACSL6*, *ANK1*, *NPR3*, *TBX20*, and *PGR*) were shared among prescapular and perirenal adipose tissues.

### Results of GO enrichment analyses

#### Enriched GO terms associated with DEGs resulting from seasonal comparisons: Finnish reindeer

Enrichment analyses based on significantly differentially expressed genes revealed several GO (Gene Ontology) terms and KEGG (Kyoto Encyclopedia of Genes and Genomes) pathways, thus highlighting the important biological and physiological activities in each tissue. From the 346 significant DEGs between FM-S and FM-W (Table 2 and S5 Table), 112 genes lacked GO annotations. Separate GO enrichment analyses were performed for the downregulated (n = 117 with GO annotation) and upregulated (n = 117 with GO annotation) genes. GO analysis indicated that 16 and 36 GO terms were significantly associated with downregulated and upregulated DEGs, respectively (S20 Table). Downregulated DEGs were represented in GO terms mainly associated with immune processes (e.g., “immune system process”, “immune response”) and stimulus (e.g., “response to stimulus”, “response to chemical”, “chemotaxis”), whereas upregulated DEGs were represented in terms associated with metabolic processes (“ATP metabolic process”, “nucleoside metabolic process”, “single-organism metabolic process”, “carbohydrate derivative metabolic process”), and transport (e.g., “proton transport”, “hydrogen transport”, “cation transmembrane transport”) in the metacarpal adipose tissue of Finnish reindeer.

In contrast to metacarpal adipose tissue, both the perirenal and prescapular adipose tissues had downregulated genes represented in a relatively higher number of GO terms (n = 29, n = 55, respectively; S21 and S22 Tables) than upregulated genes (n = 16, n = 7, respectively; S21 and S22 Tables). The DEGs upregulated during spring in Finnish reindeer perirenal adipose tissue were associated with metabolic (“cellular amide metabolic process”, “organonitrogen compound metabolic process”) and biosynthetic (“amide biosynthetic process”, “organonitrogen compound biosynthetic process”) processes, and those upregulated during winter were associated with signalling (“signalling”, “G-protein coupled receptor signalling process”, cell communication”, “single organism signalling”, “signal transduction”), and regulation (“regulation of biological process”, “regulation of cellular process”, “biological regulation”) related biological processes. Altogether, 35 of 55 GO terms associated with DEGs upregulated in the prescapular adipose tissue of Finnish reindeer from winter sampling were categorised under “biological process”, whereas none of the seven GO terms enriched in upregulated DEGs of spring samples were categorized under “biological process”. Moreover, GO terms associated with the regulation of several processes (e.g., “regulation of metabolic process,” “regulation of RNA biosynthetic process,” “regulation of biological process,” “regulation of gene expression”), response (e.g., “cellular response to organic cyclic compound”, “response to steroid hormone”, “response to lipid”, “response to hormone”, “cellular response to stimulus) and transcription activities (e.g., “transcription coactivator activity” and “transcription cofactor activity”) categorised upregulated DEGs of Finnish prescapular adipose tissue collected during winter.

#### Enriched GO terms associated with DEGs resulting from seasonal comparisons: Even reindeer

The differential expression analysis between EM-S and EM-W revealed 156 significantly DEGs (Table 2 and S8 Table), of which 55 lacked GO annotations. GO enrichment analysis of the downregulated genes (n = 77 with GO annotations) yielded seven significantly represented GO terms (S23 Table), including “metalloendopeptidase activity” and “metallopeptidase activity,” whereas no significantly enriched GO terms were found in the upregulated genes. In Even reindeer perirenal tissue, of the 103 significantly DEGs in EP-S vs EP-W (Table 2 and S9 Table), 32 genes did not have GO annotations. Gene enrichment analysis of the upregulated and downregulated DEGs revealed no statistically significantly represented GO terms. From the list of 176 significantly DEGs between ES-S and ES-W (Table 2 and S10 Table), 59 genes lacked GO annotation. Gene enrichment analysis of the upregulated and downregulated DEGs revealed no statistically significantly represented GO terms.

#### Enriched GO terms associated with DEGs resulting from location comparison

DEGs resulting from the comparison of the metacarpal adipose tissue between Even and Finnish reindeer did not reveal any GO terms. Upregulated genes in the perirenal adipose tissue of Even reindeer from spring sampling were associated only with the GO term “cofactor binding,” whereas downregulated genes were not enriched in any GO terms. Similar comparison of perirenal adipose tissue from winter sampling revealed eight and 46 GO terms associated with upregulated and downregulated genes, respectively, in Even reindeer (S24 Table).

In spring prescapular adipose tissues of Even reindeer, a total of 19 GO terms including signalling (e.g., “single organism signalling,” “signal transduction,” “G-protein coupled receptor signalling pathway”), and response to stimulus (“response to external stimulus,” “cellular response to stimulus”) were associated with upregulated genes (S25 Table), while none of the downregulated genes were enriched in any GO terms. Similar comparison of winter samples in Even reindeer revealed three overrepresented GO terms (“anion binding”, “cofactor binding” and “pyridoxal phosphate binding”) and 20 underrepresented GO terms (including six signalling terms, “G-protein coupled receptor activity”, “receptor activity” and “cell communication”) (S26 Table).

#### Enriched GO terms associated with DEGs resulting from gender comparison

From the list of 225 significantly DEGs identified in EM-F versus EM-M comparison (S17 Table), 59 genes did not have GO annotations. The GO analysis results of the upregulated (n = 58 with GO annotations) and downregulated (n = 108 with GO annotation) DEGs in female metacarpal tissue revealed one and 31 significantly represented GO terms, respectively (S27 Table). The upregulated DEGs in female metacarpal tissue were significantly enriched in only one GO term: “oxidoreductase activity, acting on paired donors, with incorporation or reduction of molecular oxygen”. The downregulated DEGs were significantly enriched in 18 biological processes, 10 molecular functions, and three cellular component categories (S27 Table). Out of the 18 biological processes, nine GO terms were associated with developmental processes, such as “circulatory system development,” “cardiovascular system development,” “blood vessel development,” “vasculature development,” and “anatomical structure development” (S27 Table) Moreover, downregulated DEGs in female metacarpal tissue were also significantly enriched in the biological process represented by the GO terms “cell adhesion,” “angiogenesis,” “blood vessel morphogenesis,” and “anatomical structure formation involved in morphogenesis” (S27 Table).

From the list of 104 significantly DEGs identified in EP-F versus EP-M comparison (S18 Table), 15 lacked GO annotations. The GO analysis results of the upregulated (n = 48 with GO annotations) DEGs in female perirenal tissue revealed six significantly represented cellular component categories, including “plasma membrane”, “cell periphery”, “plasma membrane part,” and “integral component of membrane”; no significantly represented GO terms were associated with downregulated DEGs (n = 41 with GO annotations) (S28Table).

From the list of 49 significantly DEGs identified in ES-F vs. ES-M comparison (S19 Table), seven genes lacked GO annotation. The GO analysis result of the upregulated (n = 23 with GO annotation) and downregulated (n = 19 with GO annotation) DEGs in female prescapular tissue revealed three significant represented molecular function categories and no significantly enriched GO terms, respectively. The significantly represented GO terms associated with downregulated DEGs in female reindeer prescapular tissue included “cation binding,” “ion binding,” and “metal ion binding,” indicating their role in the acquisition of mineral nutrients: iron, zinc, and calcium.

### Results of KEGG pathway analyses

#### KEGG pathways associated with seasonal comparisons

Finnish reindeer. We performed KEGG pathway analysis using GAGE to identify pathways that were differentially regulated by season (FM-S vs FM-W). Pathway enrichment analysis revealed 18 and 5 significantly downregulated and upregulated KEGG pathways, respectively, in spring metacarpal adipose tissue (S29 Table). Most of the downregulated pathways were associated with the immune system, environmental information processing, and signal transduction (S29 Table). Pathways associated with immune system included “cytokine-cytokine receptor interaction,” “chemokine signalling pathway,” “Fc gamma R-mediated phagocytosis,” “IL-17 signalling pathway,” and “T cell receptor signalling pathway” (S29 Table). In addition, the significantly downregulated pathways in spring metacarpal tissue associated with environmental information processing and signal transduction included: “TNF signalling pathway,” “MAPK signalling pathway,” “NF-kappa B signalling pathway,” “Jak-STAT signalling pathway,” and “ErbB signalling pathway” (S29 Table). Similarly, the significantly upregulated pathways in spring metacarpal tissue included “ribosome,” “oxidative phosphorylation,” “biosynthesis of secondary metabolites,” “microbial metabolism in diverse environments,” and “biosynthesis of antibiotics” (S29 Table).

KEGG pathway analysis using DEGs of FP-S vs FP-W comparison indicated that 12 and three pathways were significantly downregulated and upregulated in spring perirenal tissue, respectively (S30 Table). The downregulated pathways were mainly associated with environmental information processing and signal transduction, such as “cAMP signalling pathway,” “cGMP-PKG signalling pathway,” “Rap1 signalling pathway,” “Hippo signalling pathway,” and “MAPK signalling pathway” (S30 Table). However, pathways such as “ribosome,” “oxidative phosphorylation,” and “ribosome biogenesis in eukaryotes” (S30 Table) were significantly upregulated in spring perirenal tissue.

KEGG pathway analysis for FS-S versus FS-W DEGs revealed two and nine significantly upregulated and downregulated KEGG pathways in spring prescapular tissue, respectively (S31 Table). Pathway analysis showed a significant upregulation of two pathways, “ribosome” and “complement and coagulation cascades”. The downregulated pathways were mainly associated with environmental information processing and signal transduction: “Hippo signalling pathway–fly”, “cAMP signalling pathway”, “ErbB signalling pathway”, “MAPK signalling pathway”, “Hippo signalling pathway” and “FoxO signalling pathway”.

#### KEGG pathways associated with seasonal comparisons

Pathway analysis in Even reindeer revealed three KEGG pathways significantly upregulated in the metacarpal adipose tissues collected during spring (“ribosome”, “spliceosome,” and “ribosome biogenesis in eukaryotes”), whereas no significantly downregulated pathways were found. Analysis of DEGs from perirenal adipose tissue did not reveal any KEGG pathways. Pathways such as “complement and coagulation cascades” and “cytokine-cytokine receptor interaction” were significantly upregulated during spring in prescapular adipose tissue, while no KEGG pathways were associated with downregulated genes.

#### KEGG pathways associated with location comparisons

DEGs from the comparison of the metacarpal adipose tissue between Even and Finnish reindeer did not reveal any KEGG pathways associated with upregulated genes, whereas two KEGG pathways, “oxidative phosphorylation” and “ribosome,” were associated with genes downregulated in Even reindeer in spring (S32 Table) and a similar comparison from winter sampling revealed seven KEGG pathways, such as “ribosome biogenesis in eukaryotes”, “TNF signalling pathway”, “IL-17 signalling pathway,” “NF-kappa B signalling pathway,” and “cytokine-cytokine receptor interaction” associated with downregulated genes (S33 Table).

Upregulated genes in the perirenal adipose tissue of Even reindeer from spring sampling were associated with one KEGG pathway “ribosome,” while downregulated DEGs were not associated with any of the pathways. Similar comparison from winter sampling revealed six and 13 KEGG pathways associated with upregulated and downregulated genes, respectively, in Even reindeer (S34 Table).

While no KEGG pathways were associated with upregulated genes in prescapular adipose tissues from spring samples, two (“ribosome” and “complement and coagulation cascades”) were linked to the downregulated genes. Similar comparison in winter samples revealed 14 and one KEGG pathways associated with upregulated and downregulated genes, respectively, in Even reindeer (S35 Table).

### KEGG pathways associated with gender comparisons

KEGG pathway analysis revealed no statistically significant enriched KEGG pathways between EM-F and EM-M. “Ribosome” was the only KEGG pathway associated with DEGs from the perirenal and prescapular adipose tissues. Interestingly, in both tissues, the “ribosome” pathway was downregulated in female reindeer.

### Physiological analyses

#### Immunoblotting of UCP1 and COX4

Immunoblotting on UCP1, a protein specific to thermogenic adipocytes, was conducted on cross-section of different adipose tissues. Immunoreactivity at 32 kDa molecular weight characteristic of UCP1 was evident, although in small amounts, in the total proteins of prescapular and perirenal adipose tissues of both Even and Finnish reindeer (Fig 4). In general, prescapular adipose tissues appeared to have slightly higher UCP1 expression than perirenal adipose tissues. In Finnish reindeer, the relative expression of UCP1 was significantly higher in winter compared with spring in both prescapular (p = 0.0016*) and perirenal (p = 0.032*) adipose tissues (Fig 4c, f). The Even reindeer exhibited a similar albeit not statistically significant trend of higher UCP1 expression in winter compared with spring (Fig 4c, f).

**Fig 4.**
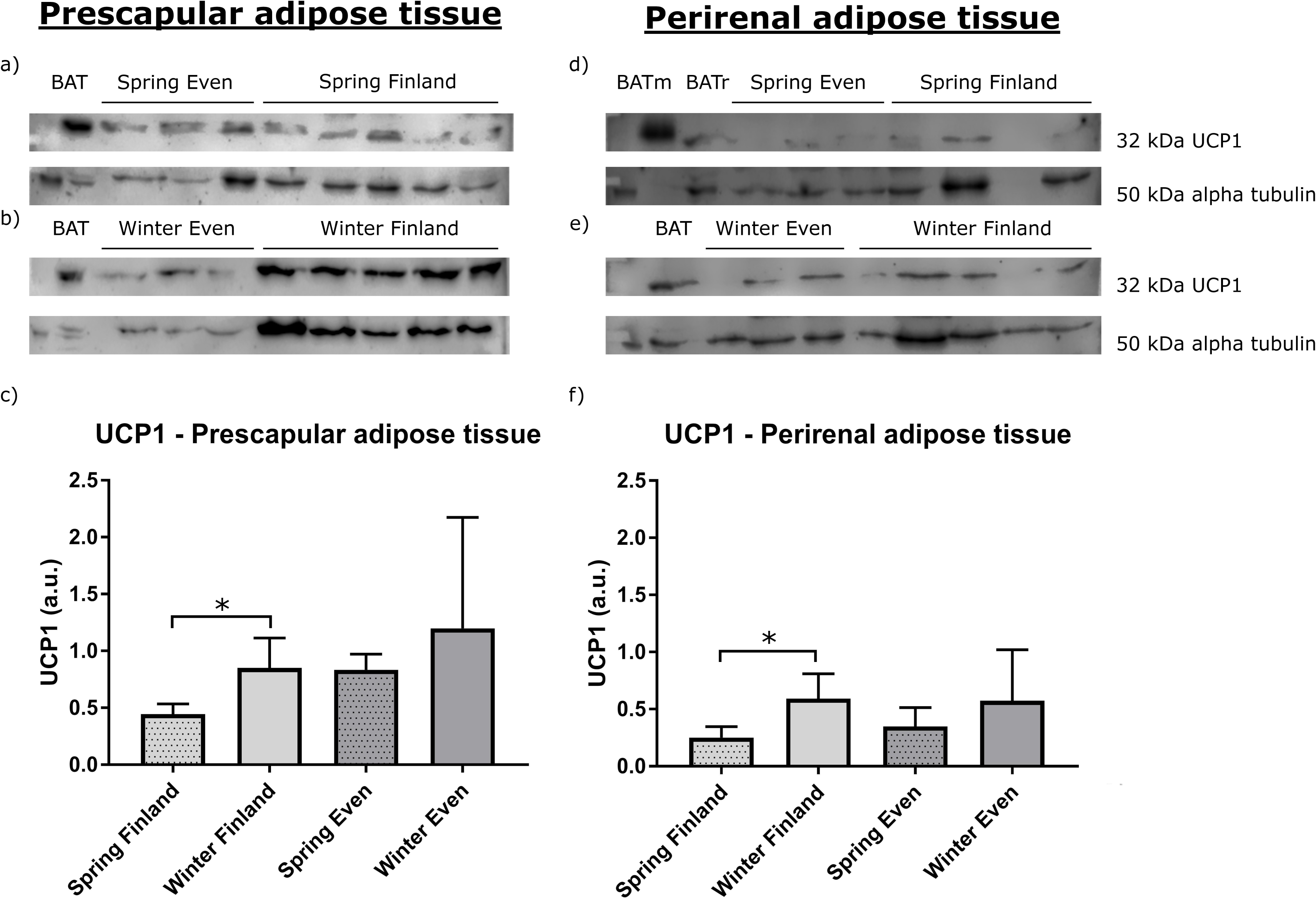
The western blots and relative expression of UCP1 from (a–c) reindeer prescapular and (d–f) perirenal adipose tissue total proteins. The upper blots on the left side show UCP1 content in prescapular adipose tissue (a) in spring in Even reindeer (n = 3) and Finnish reindeer (n = 5), (b) in winter in Even reindeer (n = 3) and Finnish reindeer (n = 5), and the lower graph shows (c) their relative expressions. The upper blots on the right side show UCP1 content in perirenal adipose tissue (d) in spring in Even reindeer (n = 3) and Finnish reindeer (n = 4), (e) in spring in Even reindeer (n = 3) and Finnish reindeer (n = 4), and the lower graph shows (f) their relative expressions. Samples contained 50 µg of total protein per lane, except the reindeer BAT (brown adipose tissue; BAT and BATr) samples with 5 µg and mouse BAT (BATm) with 1 µg of protein. Alpha tubulin was used as a loading control. The relative expression of UCP1 was normalized to alpha tubulin and presented as mean arbitrary units (a.u.) ± SD c,f). Significant differences between seasons are indicated with a bar and an asterisk (*P < 0.05).

The expression of COX4, an enzyme central to oxidative phosphorylation, was undetectable from the adipose tissue total proteins. However, we analyzed its expression in other metabolically active tissues — the liver and muscle (see Materials and methods). The relative expression level of COX4 in the liver samples was significantly higher in Finnish reindeer in winter (p = 0.008**) in comparison to spring (Fig 5). There were no statistically significant differences in muscle COX4 levels between seasons.

**Fig 5.**
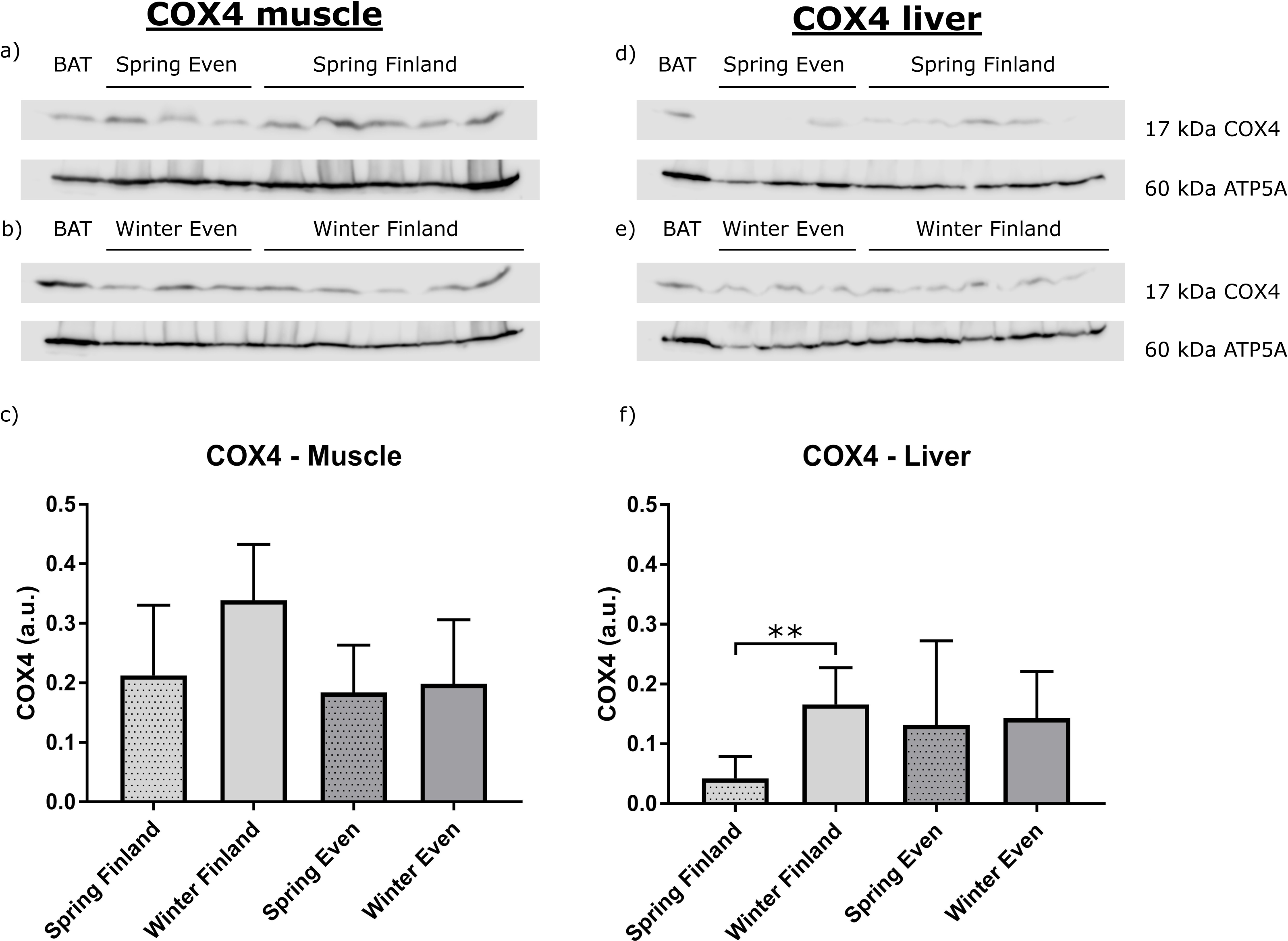
The western blots and relative expression of *COX4* from (a–c) reindeer muscle (*M. gluteobiceps femoris*) and (d–f) liver mitochondrial proteins. The upper blots on the left side show COX4 content in muscle (a) in spring in Even reindeer (n = 3) and Finnish reindeer (n = 5), (b) in winter in Even reindeer (n = 3) and Finnish reindeer (n = 5), and the lower graph shows (c) their relative expressions. The upper blots on the right side show *COX4* in liver (d) in spring in Even reindeer (n = 3) and Finnish reindeer (n = 5), e) in winter in Even reindeer (n = 3) and Finnish reindeer (n = 5), and the lower graph shows (f) their relative expressions. Samples contained 10 µg of mitochondrial protein per lane, except the reindeer BAT (brown adipose tissue) sample with 1 µg of mitochondrial protein. Mitochondrial ATP synthase (ATP5A) was used as a loading control. The relative expression of *COX4* was normalized to *ATP5A* and presented as mean arbitrary units (a.u.) ± SD c,f). Significant difference between seasons is indicated with a bar and an asterisk (**P ≤ 0.01).

#### Blood metabolites

Plasma leptin levels were at a similar range in Even and Finnish reindeer in spring, but under the detection limit (1.56 ng/ml) in both male and female Even reindeer in winter (Fig 6a). Plasma insulin levels of reindeer were at similar level in both seasons and regions (Fig 6b). Plasma HSL was significantly higher in winter in Finnish reindeer compared with spring (p = 0.016*), but there were no significant differences in the Even reindeer between seasons (Fig 6c).

**Fig 6.**
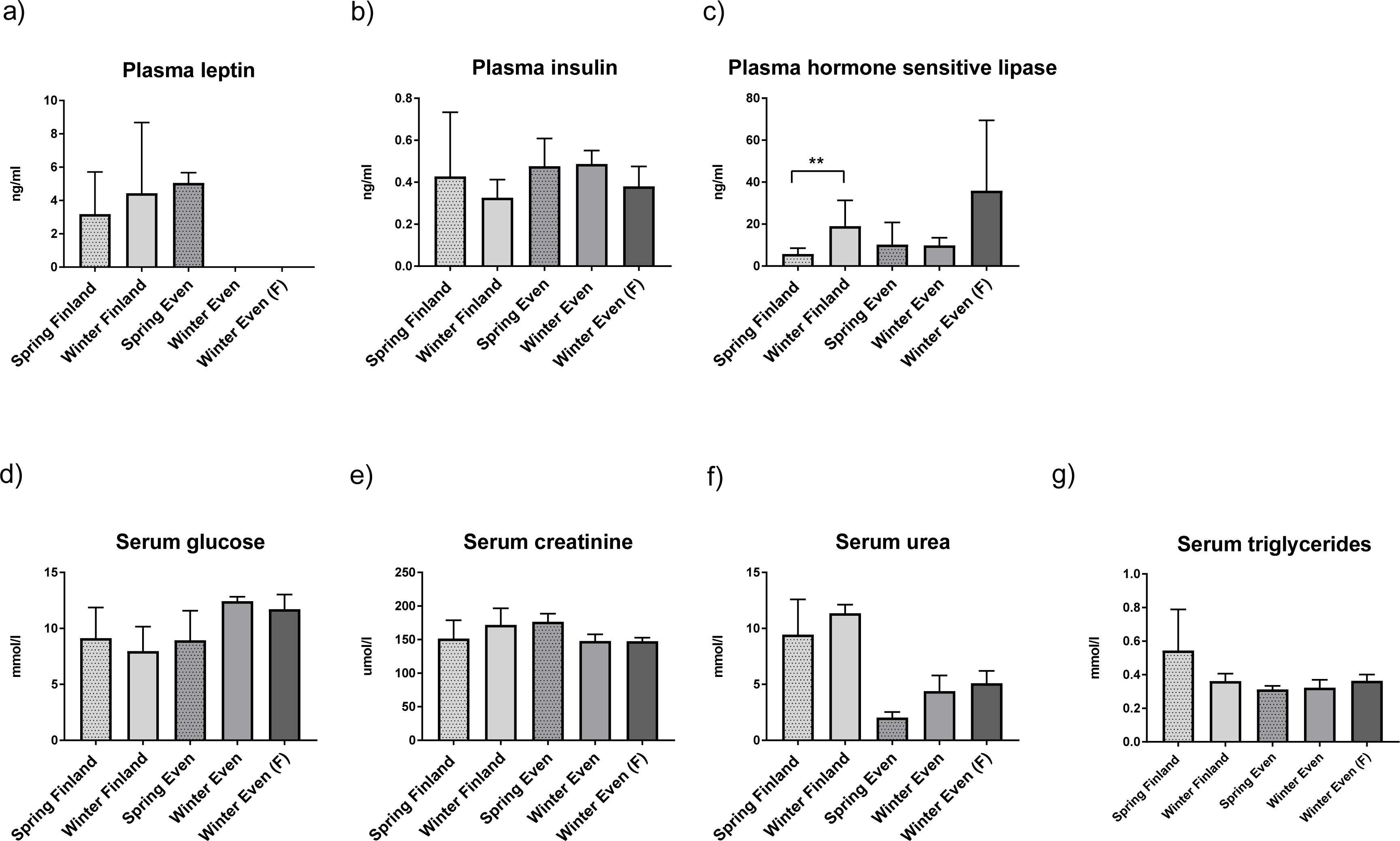
Plasma hormone and serum metabolite levels in Finnish male reindeer in winter and spring (n = 7), in male Even reindeer in winter and spring (n=3), and in Even female (F) reindeer in winter (n = 3). Plasma hormones (a) leptin, (b) insulin, (c) hormone sensitive lipase (HSL) and serum metabolites (d) glucose, (e) creatinine, (f) urea, and (g) triglyceride levels are presented as mean ± SD. Significant difference between seasons is indicated with a bar and an asterisk (**P ≤ 0.01).

Serum glucose, creatinine and triglyceride concentrations were at a similar range in both Even and Finnish reindeer in both seasons (Figures 6d, e, g). There was a trend of higher serum urea (Fig 6f) in winter samples compared to spring samples within the deer of both regions, albeit not statistically significant.

## Discussion

Adipose tissues are vital for animals living in cold environments, promoting adaptation by temperature regulation, energy homeostasis, regulation of fat deposition, and metabolism [5,6,10,11,36–40]. Here, we have investigated gene expression profiles of adipose tissues from three different anatomical depots in Finnish and Even reindeer during two seasonal time points. To the best of our knowledge, this is the first transcriptome study of adipose tissues in reindeer. Our results indicated a clear difference in gene expression profiles in metacarpal adipose tissue compared to perirenal and prescapular adipose tissues. We found that during the less optimal circumstances in early spring, mainly characterized by undernutrition, genes associated with the immune system were upregulated in perirenal and prescapular adipose tissues, while genes involved in energy metabolism were upregulated in metacarpal tissue. Interestingly, developmental, growth, and adipogenesis processes were downregulated in all three tissues during spring.

RNA-Seq is an efficient method to screen new genes, transcripts, gene expression, and differentially expressed genes in various organisms, tissues, and cells [41–43]. In this study, we identified a total of 16,362 expressed genes in focal adipose tissues, which appeared to cover approximately 60% of the list of genes available for the reindeer reference genome [3]. The present results revealed an adequate number of expressed genes in reindeer adipose tissues, which were then utilized in subsequent gene expression analysis to investigate genes associated with seasonal (early spring *vs*. early winter), location (Even reindeer *vs*. Finnish reindeer), and gender (male *vs*. female) differences.

In this study, highly abundant genes were associated with fat and lipid metabolism, thermogenesis, and energy homeostasis that are critical for the survival of reindeer during seasonal fluctuations. Several of the abundant genes involved in fat and lipid metabolism, such as *FABP4*, *FABP5*, *MT-CYB* and *ADIPOQ* (S36 Table), were highly expressed in all adipose tissues in both Finnish and Even reindeer. *Fatty acid binding protein 4* (*FABP4*) was the most highly expressed gene in all adipose tissues in both Finnish and Even reindeer. Three of the top expressed genes, including *FABP4*, *FABP5,* and *ADIPOQ,* were also found to be highly expressed in a previous adipose transcriptome profiling study in two fat-tailed sheep breeds [22]. Moreover, *FABP4* and *FABP5* were also highly expressed in the perirenal tissue of sheep [44]. Fatty acid binding proteins (FABPs) 4 and 5, encode the fatty acid binding protein found in adipocytes and epidermal cells, respectively. A previous study showed that *FABP4* and *FABP5* play an important role in thermogenesis during cold exposure and starvation [45]. Moreover, previous studies in cattle have suggested that *FABP4* plays a crucial role in fat deposition, fatty acid transport, catabolism, and metabolism [22,46,47]. Adiponectin (*ADIPOQ*), secreted by adipocytes and exclusively expressed in adipose tissues, plays an important role in modulating the regulation of fatty acid oxidation, glucose levels, and insulin sensitivity [22,48,49]. The top expressed genes associated with fat metabolism observed in the present study may play a vital role in energy homeostasis, thermoregulation, and promoting adaptation of reindeer to the challenging environment. The blood lipid, glucose, and insulin levels of reindeer support the view of homeostasis despite challenging conditions.

Tissue-wise, the gene expression profiles of metacarpal adipose tissue were remarkably different from those of prescapular and perirenal adipose tissues. The highest number of tissue-specific genes were found in metacarpal adipose tissue (Fig 1). Similarly, the PCA plot based on all expressed genes revealed a cluster for metacarpal adipose tissue, which was distinct from the samples representing other tissues (Fig 2). We found that several genes from the homeobox (HOX) family of proteins were uniquely expressed in metacarpal adipose tissue. These genes are known to play important roles in the differentiation of adipocytes [50]. Furthermore, several genes associated with cytokines and immune response appeared to be downregulated in the metacarpal tissue during spring (S5 and S8 Table). By contrast, DEGs associated with cytokines and immune response were upregulated in the perirenal and prescapular tissues during spring (S6, S7, S9 and S10 Tables). Hence, these results indicate the unique biological functions of the metacarpal tissue compared to the other two tissues.

The distinctiveness of the metacarpal adipose tissue may be due to its local niche functions in bone marrow, although it also contributes to systemic metabolism [14, 16]. Bone marrow adipose tissue has many properties in common with white adipose tissue, but it is an adipose type of its own that is currently being actively studied [14, 16]. In addition to its role as a local energy reservoir and its contribution to haematopoiesis and osteogenesis [15], metacarpal adipose tissue also secretes a variety of hormones and proteins, such as adiponectin and leptin, which have an important function in the regulation of energy metabolism [16]. The metabolic profile of metacarpal adipose tissue is composed of both white and brown fat, indicating its plasticity in performing different functions [16].

We observed more seasonal differences in the number of DEGs in the adipose tissues of Finnish reindeer (n = 1229) compared to Even reindeer (n = 386, also see Table 2). The Finnish reindeer specifically exhibited several significant DEGs associated with ATPase and ATP synthase, whereas no ATP-related genes were detected in Even reindeer. The significant DEGs associated with ATPase and ATP synthase identified in Finnish reindeer include *ATP12A, ATP5L, ATP5O, ATP6AP1L* and *ATP8* in metacarpal, *ATP1A2, ATP1B2, ATP6V0C, ATP7A* and *ATP8* in perirenal, and *ATP1A2, ATP1B2, ATP7A, ATP8B1, ATP8B3* and *ATP8* in prescapular adipose tissue. A previous study on zebrafish (*Danio rerio*) reported that feeding altered the expression of ATP-related genes [51]. The differences between Even and Finnish reindeer could be due to the influence of management (extra feeding in Finland *versus* no additional feeding in Yakutia), vegetation (relatively sparser in Yakutia), and temperature (relatively warmer in Finland). A further possible explanation for the high expression of ATP genes in Finnish reindeer is that they were fed concentrates with a high protein content (∼10% protein content) compared to their natural winter food, lichens (2–3% protein content).

In terms of seasonal comparison, we observed that the gene expression profiles of metacarpal tissue were mainly enriched for energy metabolism instead of their typical role in immune systems. Adipose tissue has been previously reported to play a role in immune and inflammatory systems [52, 53]. Earlier studies reported that adipocytes of both peripheral and bone marrow fat secrete a variety of hormones and proteins, such as pro-inflammatory and anti-inflammatory cytokines [54–56]. However, in the present Even reindeer metacarpal tissue, the downregulated genes in spring included genes associated with cytokines, such as *CCL19*, *IGDCC4*, *IL17RB*, *JCHAIN*, *LECT1* and *IGLV1-51*. Moreover, several of the downregulated DEGs in spring in Finnish and Even reindeer metacarpal adipose tissues revealed genes associated with immune system. On the other hand, the upregulated DEGs in spring in Finnish and Even reindeer metacarpal adipose tissue revealed genes associated with lipid and energy metabolism, supporting the view that bone marrow fat acts as a source of energy when reindeer are in poor condition [18, 19]. In reindeer, the proportions of unsaturated fatty acids, oleic, and linoleic acid are significantly decreased in metatarsal bone marrow fat in poor conditions in spring [19]. This may be related to their use for oxidation or other, synthetic processes. For instance, angiopoietin-like protein 8 (*ANGPTL8*) and angiopoietin-like protein 1 (*ANGPTL1*) were among the upregulated genes detected in the metacarpal tissue of Finnish and Even reindeer, respectively, in spring. *ANGPTL8* is a member of the angiopoietin-like protein (*ANGPTL*) family involved in the metabolic transition from fasting to re-feeding and plays a key role in lipid metabolism [57–59]. Depending on the location in the body, adipose tissues differ in terms of cellular composition, quantity, and proportion of adipocytes, and their capacity to produce adipocytokines [9]. During starvation, adipose tissue often limits the cytokine levels to reduce the consumption of resource/energy by the immune systems to actively decrease energy usage; subsequently, the activity of immune cells is limited [60–62]. Reindeer in both locations, and particularly Even reindeer, are in their poorest nutritional condition during spring. In this study, all cytokine genes identified in the metacarpal tissue were downregulated in spring. This might be due to the metacarpal adipose tissue being exclusively involved in thermogenesis and energy metabolism by suppressing the function of the immune system. Hence, when these reindeer experience the worst conditions during spring, the stored fat depots from metacarpal adipose tissues may be exclusively reserved for energy usage.

As revealed by the top downregulated genes in the perirenal and prescapular adipose tissues of Finnish reindeer and in all three adipose tissues of Even reindeer, biological processes associated with development, cell growth, and organogenesis were repressed during spring. The downregulation of genes involved in organogenesis, cell growth, and development in spring may indicate that the animals give less priority for growth- and development-related processes during extreme conditions; thus, the animals spend no extra energy for growth related processes. As mentioned above, during extreme conditions, the animals reduce several metabolic activities to increase the efficiency of energy usage.

Furthermore, gene expression analysis of gender differences in even reindeer adipose tissues indicated that genes associated with fatty acid metabolism and male sterility were upregulated in female and male reindeer, respectively. The upregulated genes associated with fatty acid metabolism in females include *ELOVL7* and *FABP6* in metacarpal, *ABCG1*, *ACSL6*, *PLCD3,* and *PLA2G5* in perirenal tissue, and *ACSL6* and *PLA2R1* in prescapular tissue. This result is consistent with previous studies in humans and mice [63, 64], which suggested that females show higher levels of fat deposition than males. By contrast, the upregulated genes in male reindeer revealed 10 shared genes among the three tissues, such as *UBA1*, *TXLNG*, *PRDM9*, *EIF2S3X*, *NRL*, *DDX3Y*, *KDM6A*, *ZRSR2*, *USP9X* and *ZFX*. Six of these genes, *PRDM9* [65, 66], *UBA1* [67], *EIF2S3X* [68, 69]*, DDX3Y* [70–72], *ZRSR2* [57], and *USP9X* [74], have been shown to be associated with male sterility. Moreover, it should be noted that the reindeer reference genome lacked gene annotations for the Y chromosome, and that many of the Y chromosome-specific RNA-Seq reads may have aligned to the paralog genes of the X chromosome.

We also analyzed adipose *UCP1* and *COX4* protein levels as indicators of the potential thermogenesis and metabolic state of the reindeer. We anticipated that adipose tissues of the adult reindeer are white adipose tissue but may potentially have some brown adipose tissue characteristics, considering the extreme long-term exposure of the reindeer to cold, especially in Siberia. There is no previous evidence of BAT or ‘browning’ of adipose tissues in adult reindeer, but it is well established that new-born reindeer have active BAT at birth and during their first month of life [10, 11].

Our results show the presence of *UCP1* protein in two different adipose tissues of adult reindeer, prescapular, and perirenal depots (Fig 4). Relative *UCP1* expression was significantly higher in winter compared with spring in Finnish reindeer in both prescapular and perirenal adipose tissues, likely reflecting colder weather conditions in winter. A similar trend was observed in the Even reindeer. Due to the low amount of protein, it is likely that *UCP1* does not have major thermogenic relevance. However, the findings are still interesting, as they show that *UCP1* was present in the white adipose tissues of adult reindeer. They also refer to plasticity of adipose tissue, that is, the potential to adjust its functions according to prevailing conditions [75]. In general, reindeer can cope with very low ambient temperatures in winter (-30°C) without increasing their heat production [76], and thus non-shivering thermogenesis is usually not necessary. This is mainly due to the good insulation capacity of the winter coat of reindeer, which effectively prevents heat loss. Liver *COX4* expression was significantly higher in Finnish reindeer in winter as compared to spring, and a similar trend was apparent in Even reindeer (Fig 5). This indicates an increase of oxidative phosphorylation and ATP synthesis in the liver to match the increased energy demands caused by the colder season.

Plasma leptin and insulin levels were low (Fig 6) and agree with earlier findings in reindeer [5]. Leptin is a hormone secreted by adipose tissues and plays a role in the regulation of body weight [77]. The low leptin levels suggest that the adipose tissues of reindeer were small and that the animals were striving to preserve their adipose tissues. Low leptin also agrees with adipose transcriptomics results referring to fat mobilization. Plasma hormone-sensitive lipase (HSL) was significantly higher in Finnish reindeer in the early winter than in the spring group, suggesting the mobilization of storage lipids to accommodate increased energy expenditure of male reindeer related to the breeding season.

Serum glucose, triglyceride, creatinine, and urea concentrations were similar between seasons (Fig 6), indicating that the harsh winter season is relatively well tolerated by the reindeer without severe muscle catabolism, which would be indicated by changes in the blood parameters.

### Conclusions

Collectively, our mRNA-Seq data uncovered variations in the transcriptome profiles of three adipose tissues in relation to seasonality, location, and gender differences. In general, our study showed that highly expressed genes in adipose tissues were associated with fat and lipid metabolism and thermal and energy homeostasis, promoting the adaptation of reindeer to challenging Northern Eurasian environments. Further, our results indicated a distinct gene expression profile in metacarpal adipose tissue compared to perirenal and prescapular adipose tissues. Metacarpal adipose tissue appeared to have a greater role in metabolic activities compared to the other two tissues especially during spring when the animals are experiencing the worst nutritional conditions. Moreover, reindeer from Finland and Yakutia displayed different gene expression profiles, in part owing to climatic and management differences. Thermogenic UCP1 protein was present in adipose tissues of both Even and Finnish reindeer, although in low amounts, showing that the reindeer have an option for extra heat production and thermal and energy homeostasis if needed. Taken together, the results and resources from this study will be useful for elucidating the genetics and physiology of adipose tissue for adaption to northern Eurasian conditions.

## Materials and Methods

### Sample collection for transcriptome analysis

This study includes RNA-Seq of 56 tissue samples from 19 reindeer individuals (three adult females and 16 adult males) that were randomly collected at slaughter from two different geographical regions (Inari, northern Finland and Eveno-Bytantay, Sakha, Yakutia, the Russian Federation) at two seasonal time points: winter (November–December) and spring (April) (Table 1 and S1 Table). Perirenal samples were taken from the adipose tissue around the kidneys, prescapular samples from the adipose tissue located beneath the cervical muscles in front of the scapula, and metacarpal samples from the bone marrow in the diaphysis of the metacarpal bone (left front leg). For convenience, throughout the text, the sample groups are abbreviated using reindeer location (F, E), tissue type (P, S, M), and seasonal time points (S, W). For example, FM-S represents the metacarpal tissue of Finnish reindeer collected during spring. The samples were stored in RNAlater® Solution (Ambion/QIAGEN, Valencia, CA, USA). It should be noted that three male reindeer (spring) from Yakutia were castrated, whereas those from Finland (n = 10) and autumn Yakutian males (n = 3) were uncastrated. The animals grazed on natural pastures throughout the year before the sampling. However, the Finnish reindeer were fed concentrates (Poroherkku, Raisio, Finland) for 2–8 weeks in February and March 2016 and kept in feeding pens prior to slaughter. The animals were exposed to seasonal ambient temperatures and photoperiod. The mean daily temperature in Inari, Finland varied between -16.1°C and 5.2°C before the sampling in winter (14 hours light, 10 hours dark) and between -13.2°C and 4°C before the sampling in spring (16 hours light, 8 hours dark). In northern Sakha, the daily temperature varied between -13°C and -24°C during the winter sampling (6.5 hours daylight, 17.5 hours dark) and between -9°C and - 0°C during the spring sampling (14 hours of daylight,10 hours dark). Serum and plasma samples were also collected from Sodankylä, Finland in the spring (15 hours light, 9 hours dark), where the mean daily temperatures varied between -11.4°C and 3.7°C . All protocols and sample collections were performed in accordance with the legislations approved by the Russian authorization board (FS/UVN 03/163733/07.04.2016) and the Animal Experiment Board in Finland (ESAVI/7034/04.10.07.2015).

**Table 1.** Summary of adipose tissue samples used for RNA-seq and physiological studies (the latter in brackets). Blood samples were collected from all the animals and two additional male Finnish reindeer in spring.

|  | Finnish reindeer |  | Even reindeer |  |  |
| --- | --- | --- | --- | --- | --- |
|  | Spring | Winter | Spring | Winter |  |
|  | male | male | male* | male | female |
| Metacarpal | 5 | 5 | 3 | 3 | 3 |
| Perirenal | 4 (4-5) | 5 (5) | 3 (3) | 3 (3) | 3 |
| Prescapular | 5 (5) | 5 (5) | 3 (3) | 3 (3) | 3 |

\*Castrated males (Spring samples)

### Sample collection for physiological analysis

Blood samples were taken before slaughter by a jugular venipuncture into vacuum serum and EDTA K3 tubes. The blood samples were centrifuged, and the separated serum and plasma were stored at -80°C until analysis. A total of 18 males, Finnish (n = 12) and Even (n = 6) reindeer, were examined, of which 16 were included in the RNA-seq analysis (see S1 Table). The aforementioned adipose tissues as well as additional liver and muscle (*M. gluteobiceps femoris*) samples were stored in RNAlater® solution and then used for immunoblotting analysis. We used RNAlater for preservation instead of liquid nitrogen due to the long storage of the samples in field conditions. The use of RNAlater was validated by testing both RNAlater and liquid nitrogen-preserved samples from western blotting (data not shown here).

### RNA extraction, library preparation, and sequencing

RNA extraction, library preparation, and sequencing were performed at The Finnish Functional Genomic Center (FFGC), Turku, Finland. Total RNA was extracted from adipose tissues (ca <30mg/sample) using the Qiagen AllPrep DNA/RNA/miRNA kit according to the manufacturer’s protocol. The quality of the obtained RNA was ensured with an Agilent Bioanalyzer 2100 (Agilent Technologies, Waldbronn, Germany), and the concentration of each sample was measured with a Nanodrop ND-2000 (Thermo Scientific; Wilmington, USA) and a Qbit(R) Fluorometric Quantification, Life Technologies. All samples revealed an RNA integrity number (RIN) above 7.5.

Library preparation was done according to Illumina TruSeq® Stranded mRNA Sample Preparation Guide (part #15031047). Unique Illumina TruSeq indexing adapters were ligated to each sample to pool several samples later in one flow cell lane. Library quality was inferred with an Advanced Analytical Fragment Analyzer and concentration with a Qubit fluorometer, and only good-quality libraries were sequenced.

The samples were normalized and pooled for automated cluster preparation, which was carried out with Illumina cBot station. Libraries prepared for sample YR1_SCAP_322D and FR12_SCAP_402D (S1 Table) were pooled together and run in one lane to generate a deep sequence to detect long non-coding RNAs (lncRNAS) for a future study. The remaining 61 libraries were combined in one pool and run on seven lanes of an Illumina HiSeq 3000 platform. Paired-end sequencing with 2 × 75 bp read length was used with a 8 + 8 bp dual index run. Two samples (FR9_SCAP_369D and FR13_PREN_425C) (S1 Table) suffering fed pooling error and low amounts of reads and were therefore resequenced in an extra lane. Base calling and adapter trimming were performed using Illumina’s standard bcl2fastq2 software.

### Bioinformatics analyses

The overall quality of the raw RNA-seq reads in fastq and aligned reads in BAM format were assessed using FastQC software v0.11.7 [78]. FastQC reports were summarized using MultiQC v1.7 [79]. High quality RNA-seq reads for each sample were mapped against the reindeer draft assembly [3] using Spliced Transcripts Alignment to a Reference (STAR) (version 2.6.0a) [80] with default parameters. We next generated read counts from the aligned files using the featureCounts software (version 1.6.1) from the Subread package [81] to assign reads to genes. The GTF-format annotation file associated with the reindeer draft assembly was used for gene coordinate information.

To examine the shared and uniquely expressed genes across the three adipose tissues, we used the cpm function from the edgeR library [82] to generate count-per-million (CPM) values; lowly expressed transcripts with a CPM < 0.5 were discarded.

Adipose transcriptomes are affected by the type of adipose as well as by the sex of the animal [31]. We also hypothesized that there could be differences in seasonal gene expression profiles due to changes in ambient temperature and other climatic factors and subsequent changes in body condition. Hence, we conducted differential gene expression analysis between the spring and winter sampling for each tissue and region using only male reindeer samples. In addition, we compared gene expression in samples collected from female and male Even reindeer in winter. Furthermore, to explore regional (and population) differences in gene expression, we compared expression in each tissue between E and F male reindeer. In our study, the analyzed group of animals for each tissue included at least three animals from each geographical region and season (Table 1 and S1 Table)

Raw read counts were processed using the R Bioconductor package DESeq2 [83] to perform differential gene expression and related quality control analysis. Prior to running DESeq2, lowly expressed (rowSums < 1) genes were discarded. Raw gene expression counts were normalized for differences in library size and sequencing depth using DESeq2, to enable gene expression comparisons across samples. We performed principal component analysis (PCA) to assess sample similarity using the variance stabilizing transformation (VST) method. In this study, we used fold-change and false discovery rate (FDR) filtering criteria to identify significantly differentially expressed genes (DEGs). We set absolute value of log2-fold change (LFC) to be greater than or equal to 1.5 (|log2FoldChange| > 1.5) and an adjusted p-value of 0.05 (*padj < 0.05*) to screen for significant DEGs. The Benjamini-Hochberg FDR method was used to calculate adjusted p-values.

To gain insight into the biological functions and relevance of the identified DEGs, a functional enrichment analysis was conducted using AgriGO v2.0 [84]. In the AgriGO analysis toolkit, to detect the significantly enriched GO terms, default parameters were used in the “Advanced options”: Fisher as the statistical test method, Yekutieli for multiple test correction at a significance level threshold 0.05 (FDR < 0.05), and minimum number of mapping entries 5. In this analysis, the GO annotation file from the *de novo* assembled reindeer genome [3] was used as a background reference. Furthermore, to explore the biological pathways associated with the DEGs, we performed Kyoto Encyclopaedia of Genes and Genomes (KEGG) pathway analysis using the GAGE [85] Bioconductor package. The significantly enriched pathways were identified based on the q-values obtained from a Fisher’s exact test (*q-value < 0.1*).

### Physiological analyses

#### Immunoblotting analysis of proteins

Adipose tissue samples were homogenized and dissolved in lysis buffer (25 mM Tris [pH 7.4], 0.1 mM EDTA, 1 mM DTT, 15µl/ml protease inhibitor cocktail (Sigma, St Luis, MO, USA) to extract total protein content. Insoluble material was removed from the extracts by centrifugation (13,000 g, 10 min, +4°C). Mitochondrial proteins were extracted from reindeer muscle, liver, reindeer calf prescapular brown adipose tissue (BAT), and mouse BAT samples as described previously [86]. Both total and mitochondrial protein concentrations were determined using the Bradford method (Bio-Rad protein assay, Bio-Rad Laboratories GmbH, München, Germany). Protein extract volumes equivalent to 50–75 μg of total protein were concentrated into smaller volumes by lyophilizing the samples with a Savant Speed Vac Plus SC210A centrifugal evaporator (Thermo Fisher Scientific, Rockford, USA) in cooled conditions for 1 hour. Due to the low number of mitochondria in adipose tissue, total protein extractions were used for the immunoblotting of UCP1.

The proteins were separated electrophoretically using a 4–12% gradient gel and transferred to a nitrocellulose membrane (Bio-Rad, Trans-Blot® Transfer Medium, Pure Nitrocellulose Membrane [0.2 μ CA, USA). The membranes were incubated overnight with UCP1 antibody and loading control protein alpha tubulin antibody for the total protein adipose tissue samples (1:1.000 UCP1 Polyclonal Antibody, cat no. PA1-24894, 1:500 alpha Tubulin Polyclonal Antibody, cat no. PA5-16891, Invitrogen, Thermo Fisher Scientific, Rockford, USA). Membranes with liver and muscle mitochondrial protein samples were incubated overnight with COX4 antibody (1:1.000 COX4 Polyclonal Antibody, cat no. PA5-17511, Thermo Fisher Scientific, Rockford, USA) and loading control protein ATP5A antibody (1:1.000 anti-ATP5A antibody, cat no. ab151229, Abcam, Cambridge, UK). After the primary antibody treatments, the membranes were incubated with a secondary antibody (1:25.000, Goat anti-Rabbit IgG (H+L) horseradish peroxidase conjugate, cat no. 31460, Invitrogen, Thermo Fisher Scientific, Rockford, USA) for 1 hour. Chemiluminescence for UCP1 and COX4 was detected with SuperSignal West Femto Maximum Sensitivity Substrate (cat no. 34095, Thermo Fisher Scientific, Rockford, USA) according to the manufacturer’s instructions. Blots were visualized with Odyssey® Fc imaging system (LI-COR Biosciences, Ltd, Cambridge, UK). Positive immunoreactivity for UCP1 with mouse BAT mitochondria and with prescapular BAT mitochondria from newborn reindeer were used as reference samples. Results were normalized with the loading control optical density for alpha tubulin and ATP5A for UCP1 and COX4, respectively.

#### Blood metabolites

Plasma leptin concentration was assayed using a multispecies leptin RIA kit (Cat#XL-85K, Millipore, Billerica, Massachusetts, USA). The validity test for reindeer showed a linear correlation between the label and sample concentration. A sensitive bovine, ovine, rat, and mouse insulin RIA kit (Cat#SRI-13K, Millipore, Billerica, Massachusetts, USA) was used to measure plasma insulin. Plasma hormone-sensitive lipase levels (HSL) were estimated by bovine hormone sensitive ELISA Kit (Cat#MBS033124, MyBioSource, San Diego, CA, USA). Serum glucose, triglyceride, creatinine, and urea concentrations were determined with enzymatic colorimetric analyses in NordLab, Oulu, Finland.

#### Statistical analyses for immunoblotting and blood metabolite analyses

Statistical analysis for the multiple comparisons was performed with an independent-samples Kruskal-Wallis test followed by an independent samples Mann-Whitney U-test. Significance values were adjusted with the Bonferroni correction for multiple tests. Statistical analyses were performed using the IBM SPSS Statistics 21 Data Editor software (IBM, Armonk, NY, USA). P-values below 0.05 were considered statistically significant. The results of the relative peptide expressions are presented as the mean ± SD.

## Supporting information

**S1 Fig.** PCA plot based on the region for each tissue. (A) metacarpal, (B) perirenal, and (C) prescapular.

**S2 Fig.** Heatmap plot of the top 25 genes with the highest genetic variance across all samples.

**S3 Fig.** The number of shared and uniquely significant DEGs between three adipose tissues in Finnish reindeer due to seasonal differences. Significant DEGs detected in Finnish reindeer for three adipose tissues due to seasonal change: FM-S *vs.* FM-W, FP-S *vs.* FP-W, and FS-S *vs.* FS-W.

**S4 Fig.** Volcano plot of differentially expressed genes between spring and winter for metacarpal adipose tissue in Finnish reindeer (FM-S *vs.* FM-W).

**S5 Fig.** Volcano plot of differentially expressed genes between spring and winter for perirenal adipose tissue in Finnish reindeer (FP-S *vs.* FP-W).

**S6 Fig.** Volcano plot of differentially expressed genes between spring and winter for prescapular adipose tissue in Finnish reindeer (FS-S *vs.* FS-W).

**S7 Fig.** The number of shared and unique significant DEGs between three adipose tissues in Even reindeer due to seasonal differences. Significant DEGs detected in Even reindeer for three adipose tissues due to seasonal change (EM-S *vs.* EM-W, EP-S *vs.* EP-W and ES-S *vs.* ES-W).

**S8 Fig.** Volcano plot of differentially expressed genes between early spring and early winter for metacarpal adipose tissue in Even reindeer (EM-S *vs.* EM-W).

**S9 Fig.** Volcano plot of differentially expressed genes between early spring and early winter for metacarpal adipose tissue in Even reindeer (EM-S *vs.* EM-W).

**S10 Fig.** Volcano plot of differentially expressed genes between early spring and early winter for metacarpal adipose tissue in Even reindeer (EM-S *vs.* EM-W).

**S11 Fig.** The number of shared and unique significant DEGs in the three adipose tissues detected between Even reindeer and Finnish reindeer due to regional differences in early spring. Significant DEGs detected in three adipose tissues due to regional differences in early spring: EM-S *vs.* EM-S, EP-S *vs.* EP-S and ES-S *vs.* ES-S.

**S12 Fig.** The number of shared and unique significant DEGs in the three adipose tissues detected between Even reindeer and Finnish reindeer due to regional differences in early winter. Significant DEGs detected in three adipose tissues due to regional differences in early winter: EM-W *vs.* EM-W, EP-W *vs.* EP-W and ES-W *vs.* ES-W.

**S1 Table.** Statistics of clean data.

**S2 Table.** STAR mapping statistics.

**S3 Table.** Summary of expressed genes in each tissue (sheet). List of expressed genes in Finnish reindeer metacarpal adipose tissue in spring (FM-S) (sheet 2). List of expressed genes in Finnish reindeer perirenal adipose tissue in spring (FP-S) (sheet 3). List of expressed genes in Finnish reindeer prescapular adipose tissue in spring (FS-S) (sheet 4). List of expressed genes in Finnish reindeer metacarpal adipose tissue in winter (FM-W) (sheet 5). List of expressed genes in Finnish reindeer perirenal adipose tissue in winter (FP-W) (sheet 6). List of expressed genes in Finnish reindeer prescapular adipose tissue in winter (FS-W) (sheet 7). List of expressed genes in Even reindeer metacarpal adipose tissue in spring (EM-S) (sheet 8). List of expressed genes in Even reindeer perirenal adipose tissue in spring (EP-S) (sheet 9). List of expressed genes in Even reindeer prescapular adipose tissue in spring (ES-S) (sheet 10). List of expressed genes in Even reindeer metacarpal adipose tissue in winter (EM-W) (sheet 11). List of expressed genes in Even reindeer perirenal adipose tissue in winter (EP-W) (sheet 12). List of expressed genes in Even reindeer prescapular adipose tissue in winter (ES-W) (sheet 13).

**S4 Table.** Uniquely expressed genes in metacarpal adipose tissue shared by Finnish and Even reindeer in both seasons.

**S5 Table.** Significantly differentially expressed genes between spring and winter in Finnish reindeer metacarpal tissue (FM-S *vs.* FM-W).

**S6 Table.** Significantly differentially expressed genes between spring and winter in Finnish reindeer perirenal tissue (FP-S *vs.* FP-W).

**S7 Table.** Significantly differentially expressed genes between spring and winter in Finnish reindeer prescapular tissue (FS-S vs. FS-W).

**S8 Table.** Significantly differentially expressed genes between spring and winter in Even reindeer metacarpal tissue (EM-S *vs.* EM-W).

**S9 Table.** Significantly differentially expressed genes between spring and winter in Even reindeer perirenal tissue (EP-S *vs*. EP-W).

**S10 Table.** Significantly differentially expressed genes between spring and winter in Even reindeer prescapular tissue (ES-S *vs.* ES-W).

**S11 Table.** Significantly differentially expressed genes between Even and Finnish reindeer in metacarpal adipose tissue in spring (EM-S *vs.* FM-S).

**S12 Table.** Significantly differentially expressed genes between Even and Finnish reindeer in perirenal adipose tissue in spring (EP-S *vs.* FM-S).

**S13 Table.** Significantly differentially expressed genes between Even and Finnish reindeer in prescapular adipose tissue in spring (ES-S *vs.* FS-S).

**S14 Table.** Significantly differentially expressed genes between Even and Finnish reindeer in metacarpal adipose tissue in winter (EM-W *vs.* FM-W).

**S15 Table.** Significantly differentially expressed genes between Even and Finnish reindeer in perirenal adipose tissue in winter (EP-W *vs.* FPW).

**S16 Table.** Significantly differentially expressed genes between Even and Finnish reindeer in prescapular adipose tissue in winter (ES-W *vs.* FS-W).

**S17 Table.** Significantly differentially expressed genes between female and male Even reindeer metacarpal adipose tissue (EM-F *vs.* EM-M).

**S18 Table.** Significantly differentially expressed genes between female and male Even reindeer perirenal adipose tissue (EP-F *vs.* EP-M).

**S19 Table.** Significantly differentially expressed genes between female and male Even reindeer prescapular adipose tissue (ES-F *vs.* ES-M).

**S20 Table.** List of significantly enriched GO terms associated with significantly downregulated DEGs (sheet 1) and upregulated DEGs (sheet 2) in Finnish reindeer metacarpal adipose tissue in spring compared to winter.

**S21 Table.** List of significantly enriched GO terms associated with significantly downregulated DEGs (sheet 1) and upregulated DEGs (sheet 2) in Finnish reindeer perirenal adipose tissue in spring compared to winter.

**S22 Table.** List of significantly enriched GO terms associated with significantly downregulated DEGs (sheet 1) and upregulated DEGs (sheet 2) in Finnish reindeer prescapular adipose tissue in spring compared to winter.

**S23 Table.** List of significantly enriched GO terms associated with significantly downregulated DEGs in Even reindeer metacarpal adipose tissue in spring compared to winter.

**S24 Table.** List of significantly enriched GO terms associated with significantly downregulated DEGs (sheet 1) and upregulated DEGs (sheet 2) in Even reindeer perirenal adipose tissue compared to Finnish reindeer perirenal adipose tissue in winter.

**S25 Table.** List of significantly enriched GO terms associated with significantly upregulated DEGs in Even reindeer prescapular adipose tissue compared to Finnish reindeer prescapular adipose tissue in spring.

**S26 Table.** List of significantly enriched GO terms associated with significantly downregulated DEGs (sheet 1) and upregulated DEGs (sheet 2) in Even reindeer prescapular adipose tissue compared to Finnish reindeer prescapular adipose tissue in winter.

**S27 Table.** List of significantly enriched GO terms associated with significantly downregulated DEGs in female Even reindeer metacarpal adipose tissue compared to male Even reindeer metacarpal adipose tissue.

**S28 Table.** List of significantly enriched GO terms associated with significantly upregulated DEGs in female Even reindeer perirenal adipose tissue compared to male Even reindeer perirenal adipose tissue.

**S29 Table.** List of significantly enriched KEGG pathways associated with significantly downregulated DEGs (sheet 1) and upregulated DEGs (sheet 2) in Finnish reindeer metacarpal adipose tissue in spring compared to winter.

**S30 Table.** List of significantly enriched KEGG pathways associated with significantly downregulated DEGs (sheet 1) and upregulated DEGs (sheet 2) in Finnish reindeer perirenal adipose tissue in spring compared to winter.

**S31 Table.** List of significantly enriched KEGG pathways associated with significantly downregulated DEGs (sheet 1) and upregulated DEGs (sheet 2) in Finnish reindeer prescapular adipose tissue in spring compared to winter.

**S32 Table.** List of significantly enriched KEGG pathways associated with significantly downregulated DEGs in Even reindeer metacarpal adipose tissue compared to Finnish reindeer metacarpal adipose tissue in spring.

**S33 Table.** List of significantly enriched KEGG pathway associated with significantly downregulated DEGs in Even reindeer metacarpal adipose tissue compared to Finnish reindeer metacarpal adipose tissue in winter.

**S34 Table.** List of significantly enriched KEGG pathways associated with significantly downregulated DEGs (sheet 1) and upregulated DEGs (sheet 2) in Even reindeer perirenal adipose tissue compared to Finnish reindeer perirenal adipose tissue in winter.

**S35 Table.** List of significantly enriched KEGG pathways associated with significantly downregulated DEGs (sheet 1) and upregulated DEGs (sheet 2) in Even reindeer prescapular adipose tissue compared to Finnish reindeer prescapular adipose tissue in winter.

**S36 Table.** List of the top 25 most abundant genes expressed in the three adipose tissues based on mean TPM in Finnish reindeer (sheet1) and Even reindeer (sheet2).

## Author contributions

**Conceptualization:** Juha Kantanen

**Data curation:** Melak Weldenegodguad, Kisun Pokharel, Laura Niiranen

**Formal analysis:** Melak Weldenegodguad, Laura Niiranen

**Funding acquisition:** Juha Kantanen

**Investigation:** Melak Weldenegodguad, Kisun Pokharel, Laura Niiranen, Päivi Soppela

**Methodology:** Melak Weldenegodguad, Kisun Pokharel, Laura Niiranen, Päivi Soppela

**Project Administration:** Juha Kantanen

**Resources:** Päivi Soppela, Innokentyi Ammosov, Mervi Honkatukia, Heli Lindeberg, Jaana Peippo, Tiina Reilas, Nuccio Mazzullo, Florian Stammler, Juha Kantanen

**Supervision:** Juha Kantanen, Tommi Nyman, Arja Tervahauta, Päivi Soppela **Visualization:** Melak Weldenegodguad, Kisun Pokharel, Laura Niiranen, Tommi Nyman

**Writing – original draft preparation:** Melak Weldenegodguad, Kisun Pokharel, Laura Niiranen, Juha Kantanen

**Writing – review and editing:** Melak Weldenegodguad, Kisun Pokharel, Laura Niiranen, Päivi Soppela, Innokentyi Ammosov, Mervi Honkatukia, Heli Lindeberg, Jaana Peippo, Tiina Reilas, Nuccio Mazzullo, Kari A. Mäkelä, Tommi Nyman, Arja Tervahauta, Karl-Heinz Herzig, Florian Stammler, Juha Kantanen

### Funding disclosure

This study was funded by the Academy of Finland in the Arctic Research Programme ARKTIKO (decision number 286040).

## Acknowledgements

We thank reindeer herders from the Sallivaara and Oraniemi reindeer herding districts in Finland and from the Eveno-Bytantay District in Sakha, Russia for providing reindeer for our research and for helping with the field work. We thank Tuula-Marjatta Hamama for RNA extractions. We thank the CSC-IT Center for Science, Finland, for computational resources, and the Finnish Functional Genomics Centre supported by the University of Turku, Åbo Akademi University, and Biocenter Finland for RNA sequencing.

