## Supplementary figures and images for "Adipose gene expression profiles reveal novel insights into the adaptation of northern Eurasian semi-domestic reindeer (*Rangifer tarandus*)"

### Figure_S1.pdf

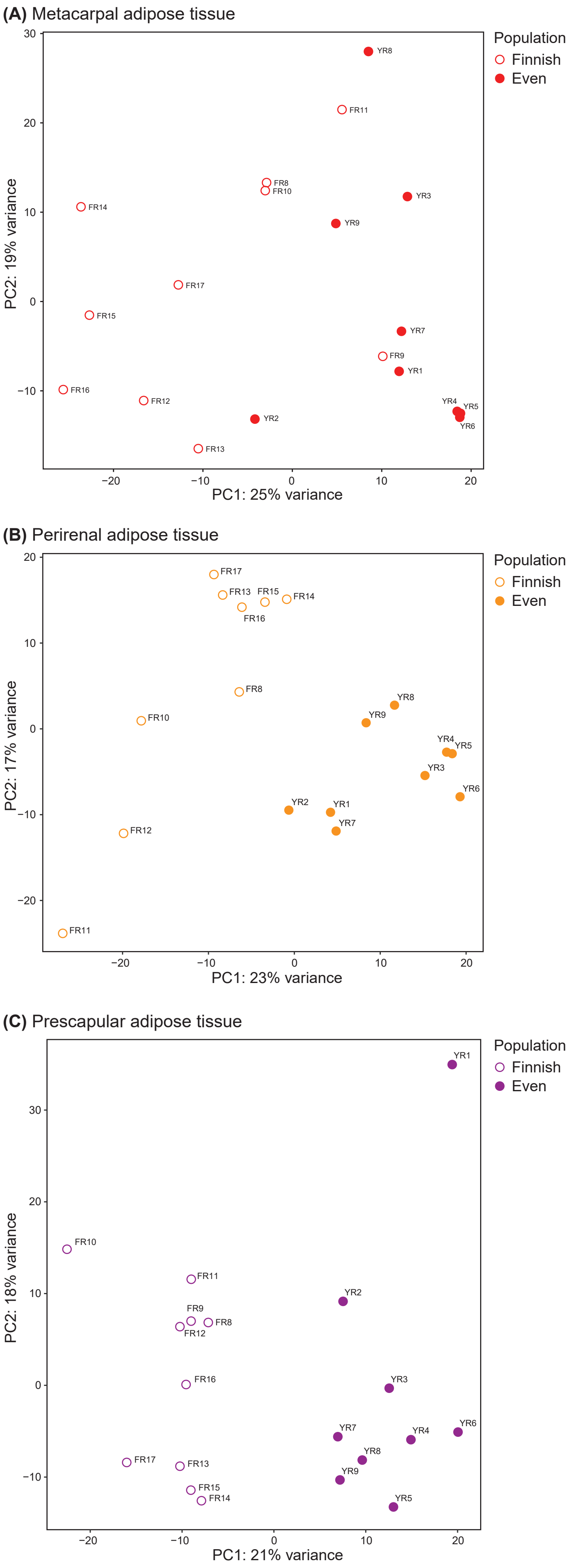

### Figure_S2.pdf

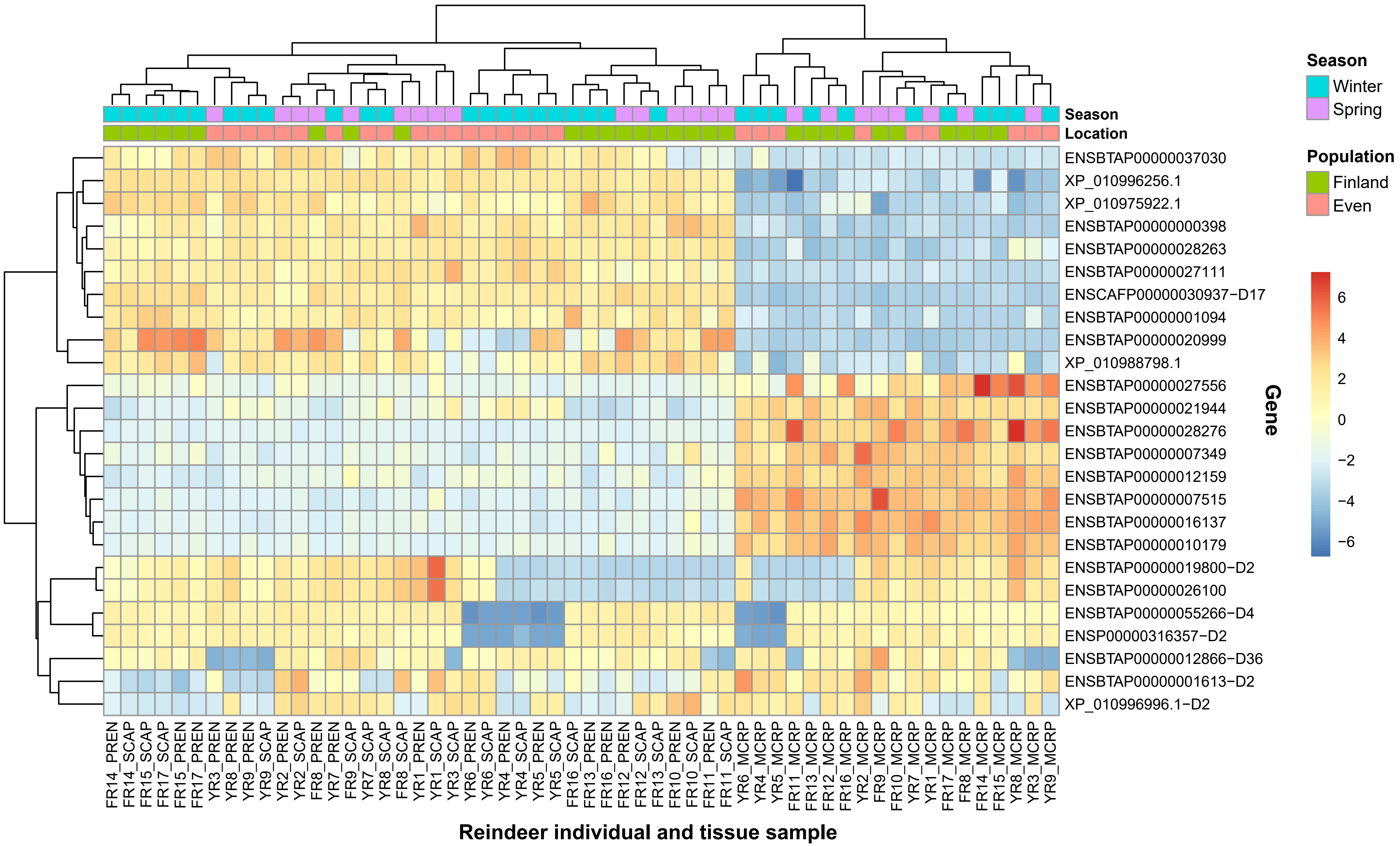

### Figure_S3.pdf

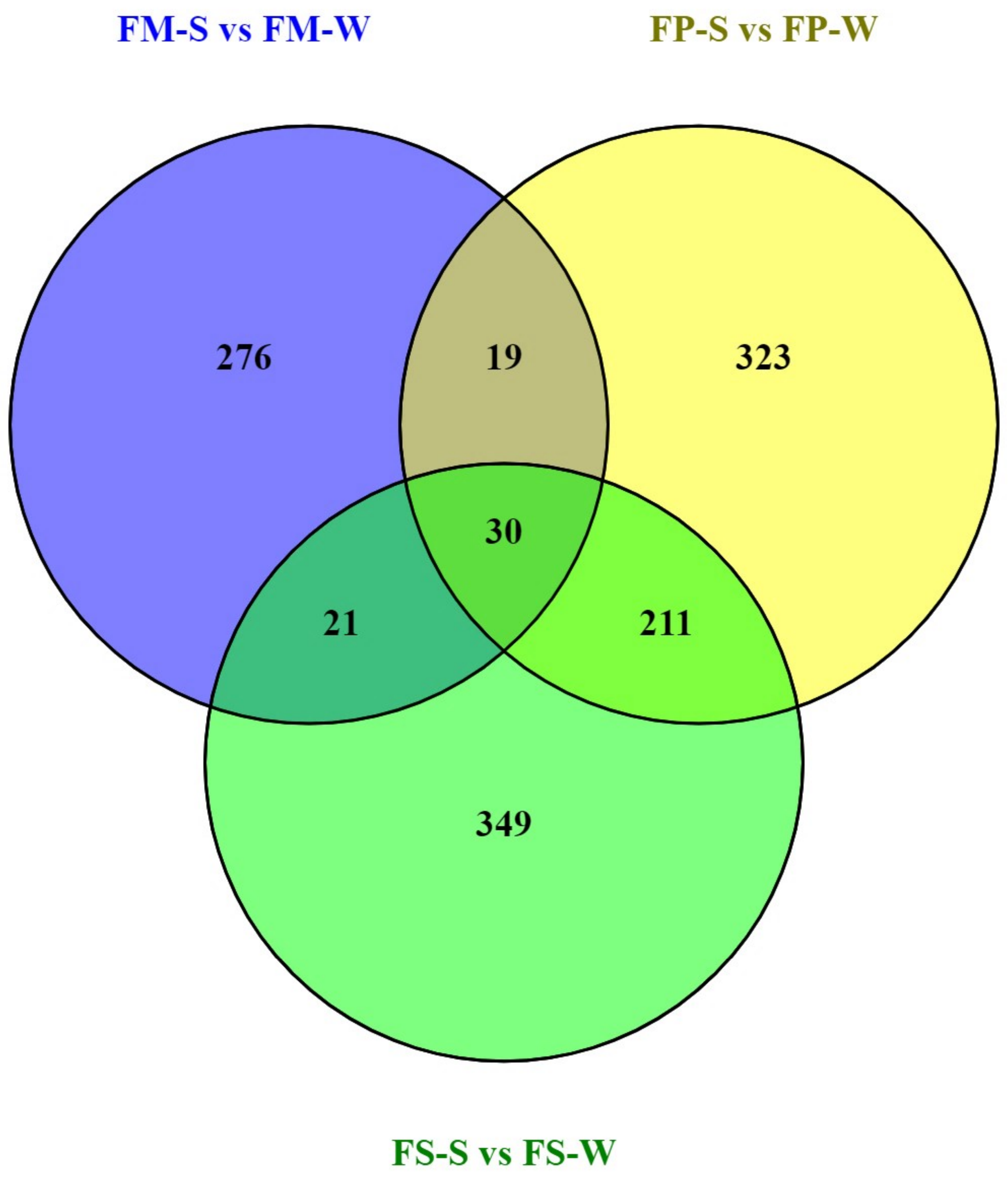

### Figure_S4.pdf

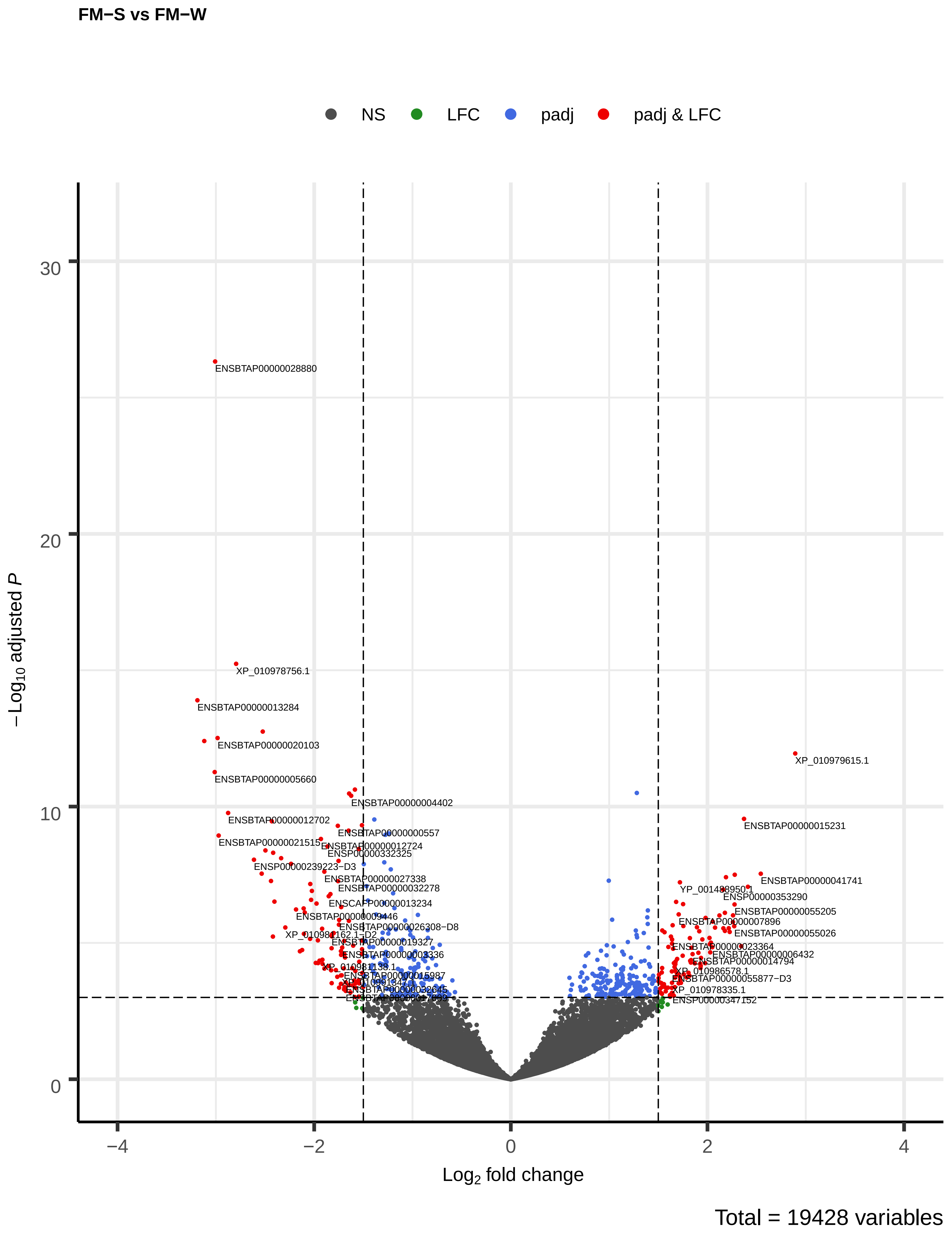

### Figure_S5.pdf

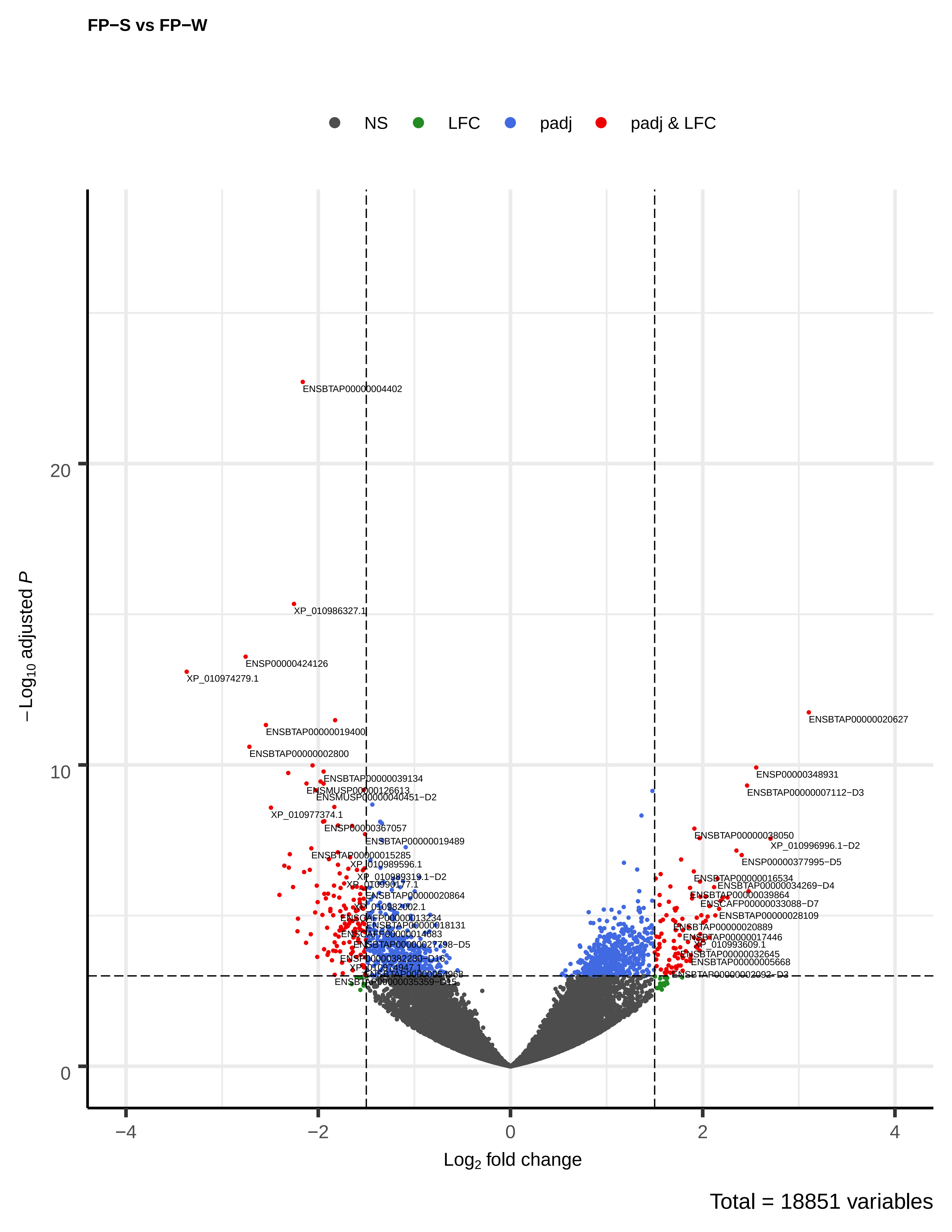

### Figure_S6.pdf

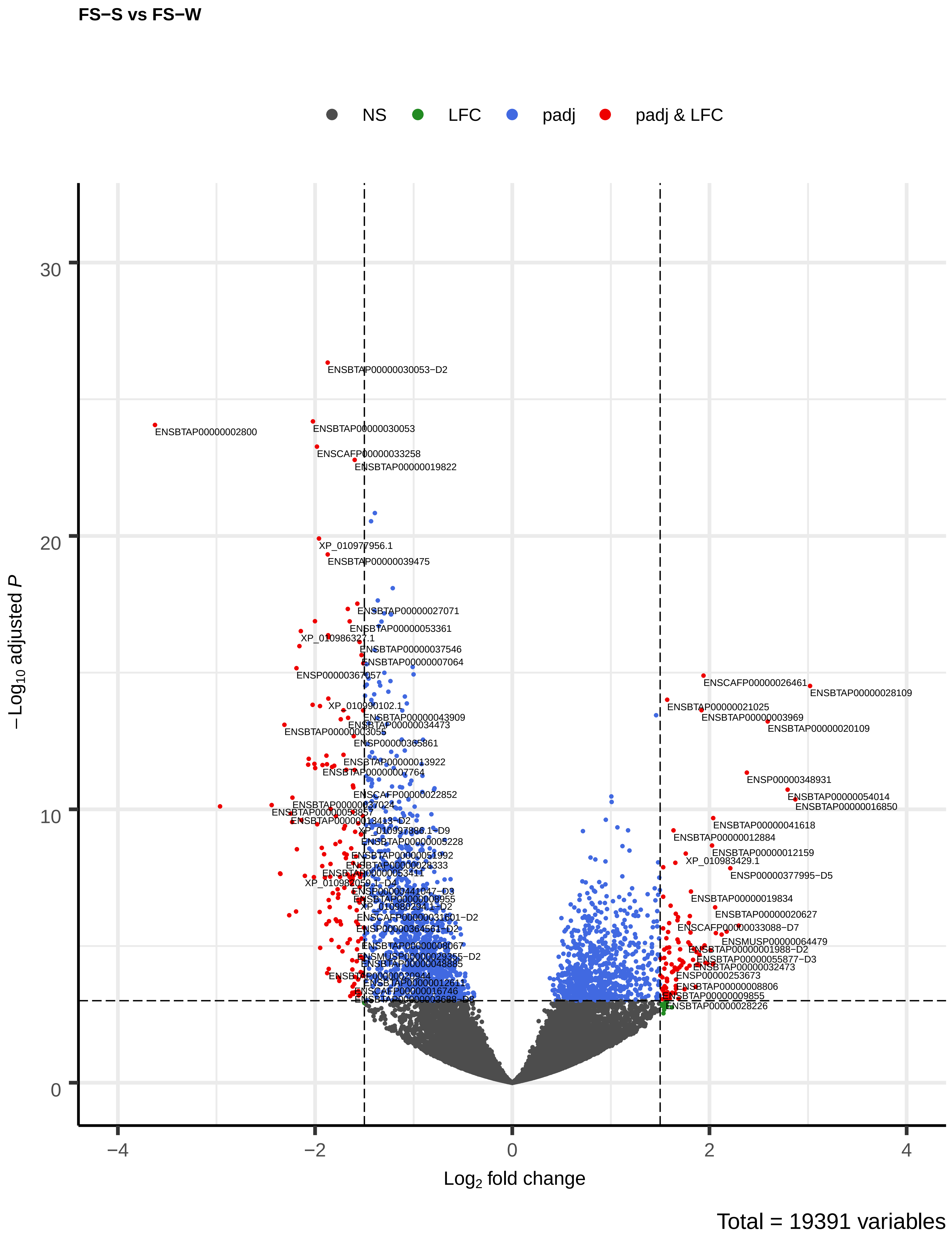

### Figure_S7.pdf

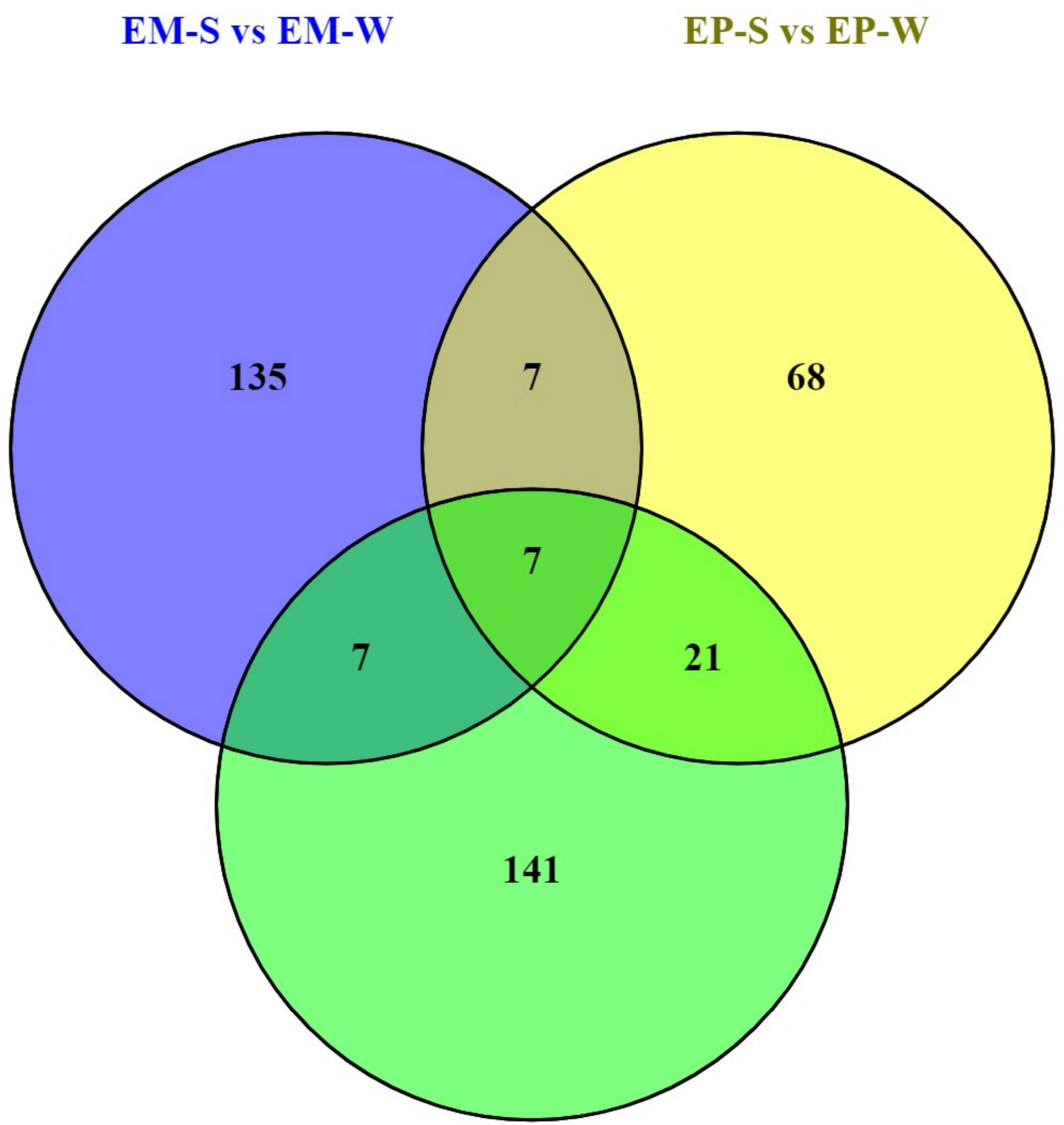

### Figure_S8.pdf

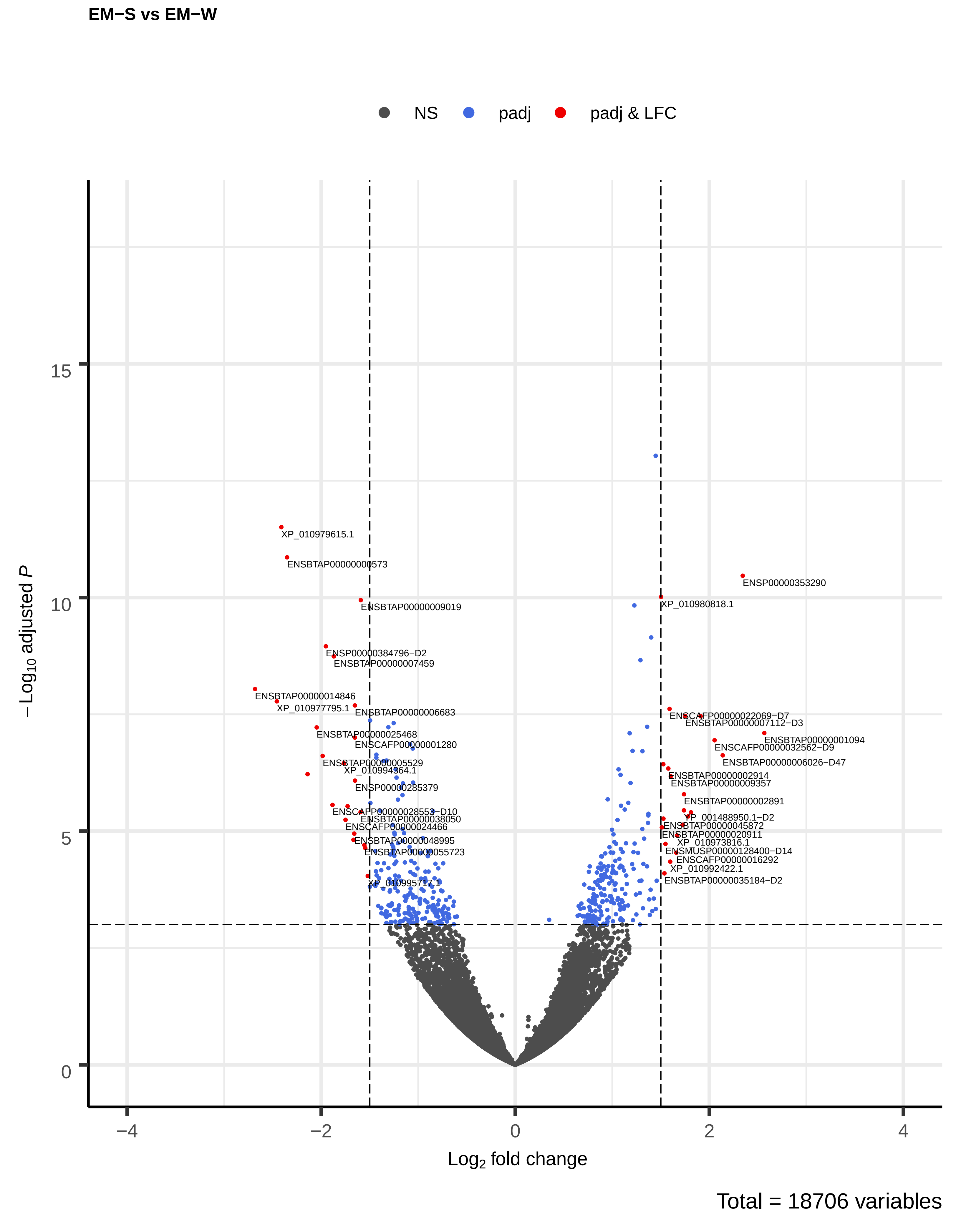

### Figure_S9.pdf

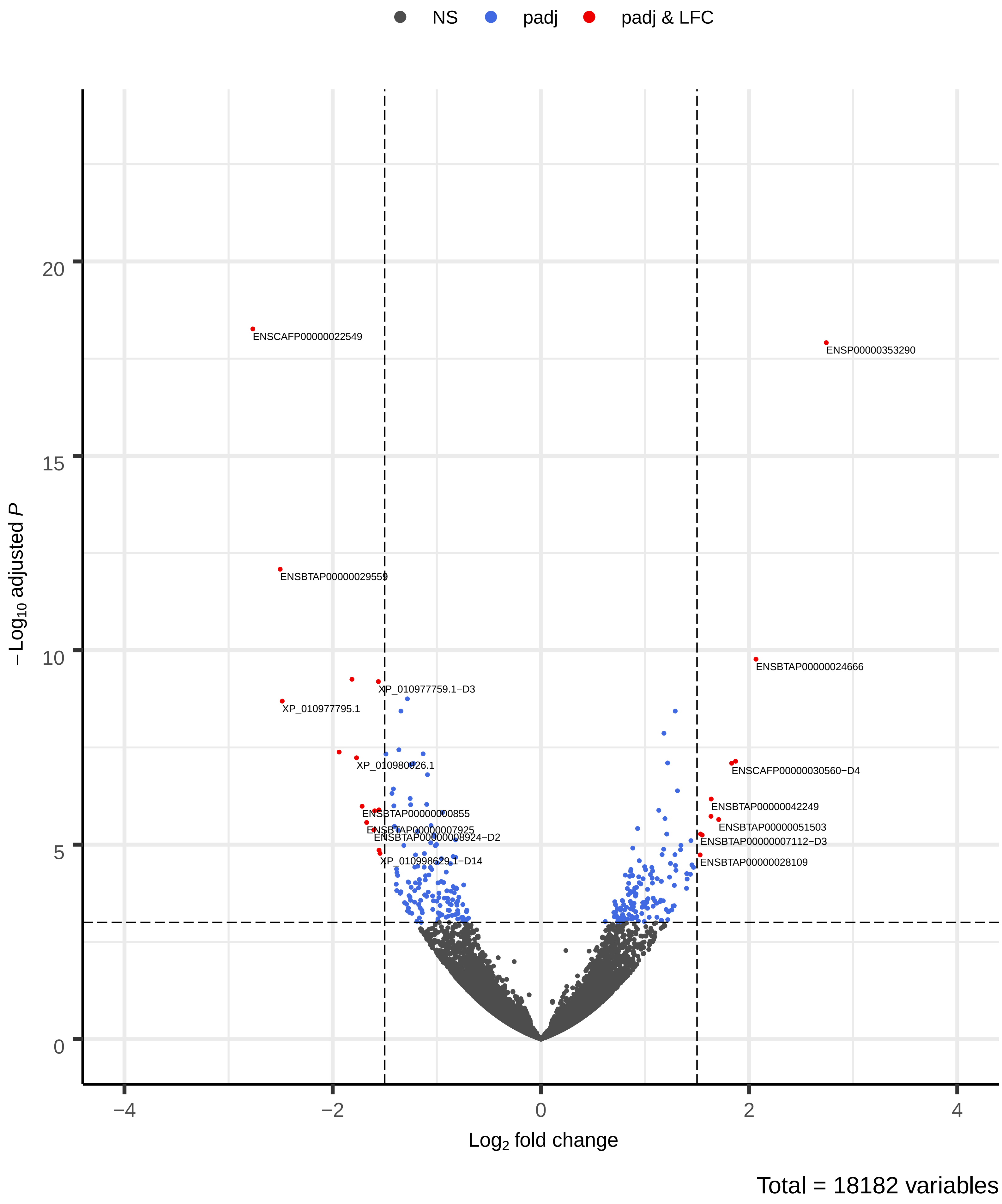

### Figure_S10.pdf

**ES−S vs ES−W**

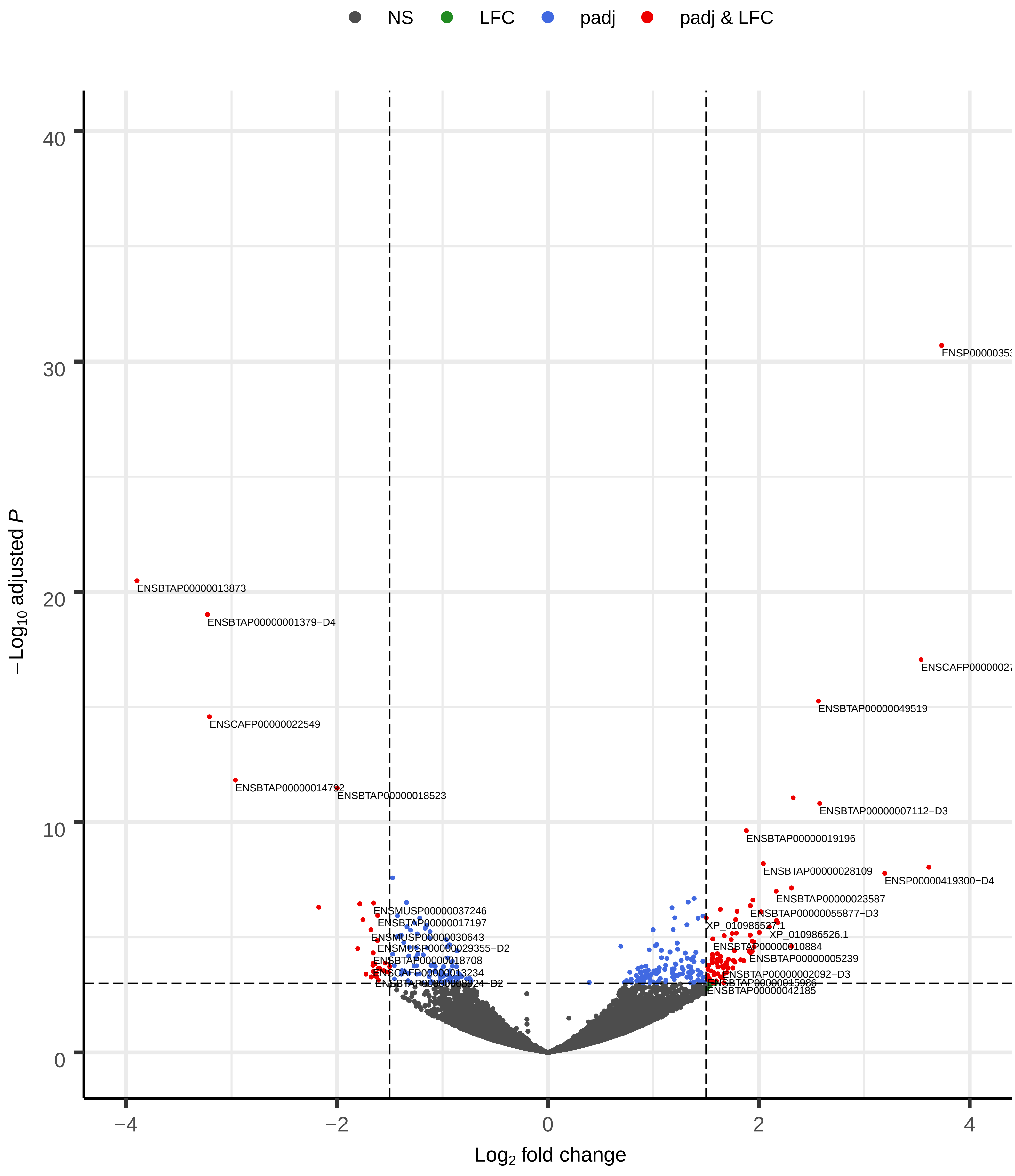


Total = 18707 variables

### Figure_S11.pdf

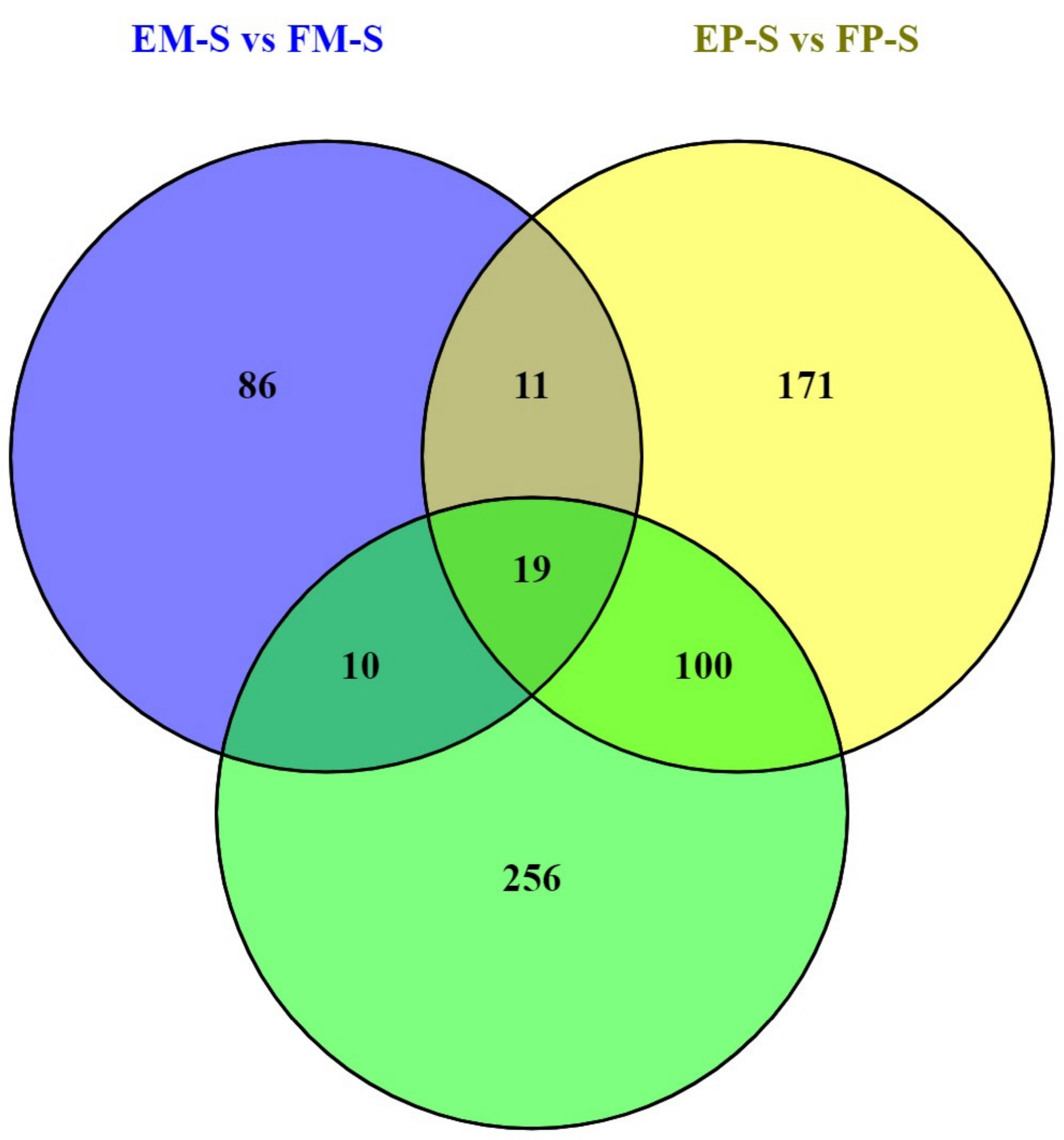

### Figure_S12.pdf

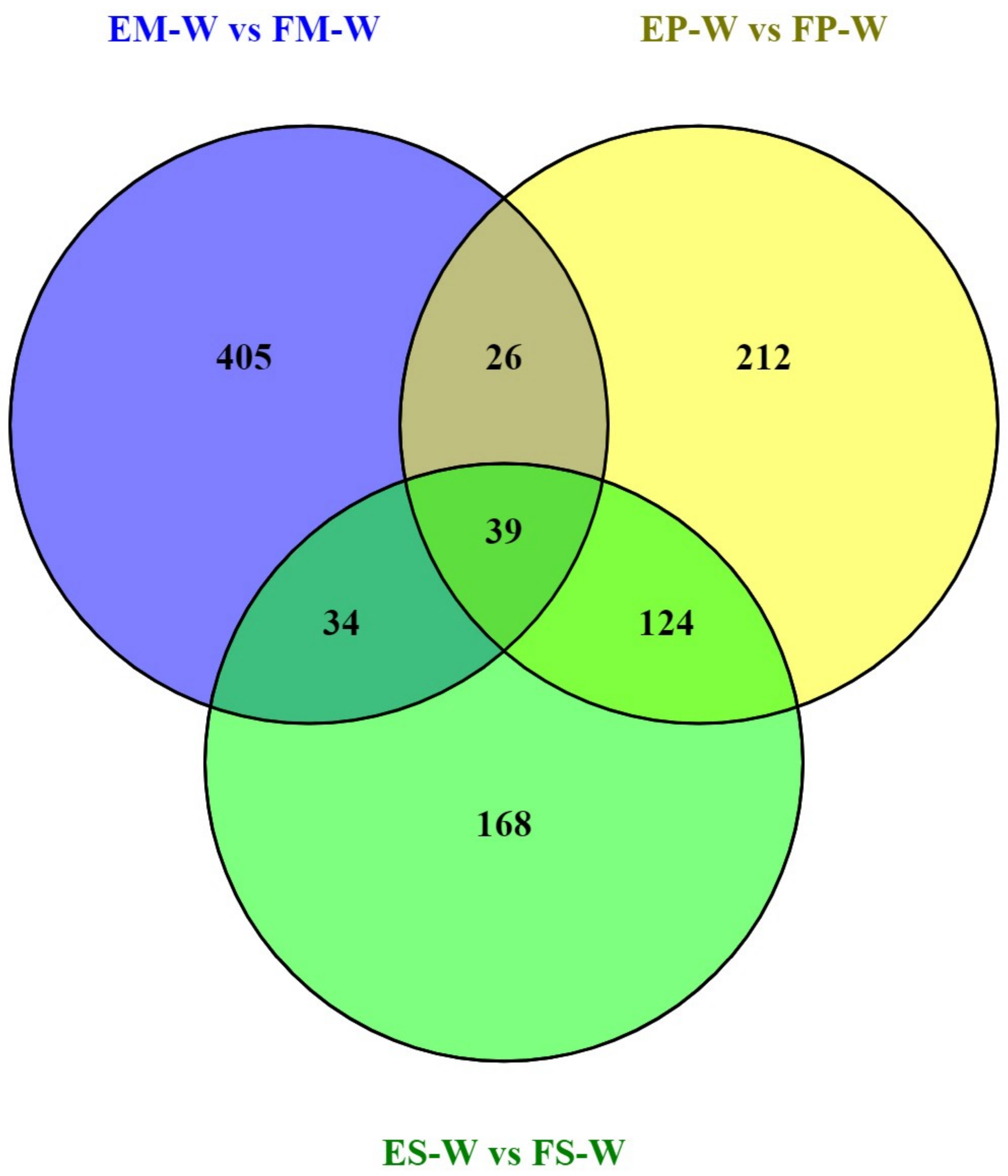
