## Supplementary material for "Adipose gene expression profiles reveal novel insights into the adaptation of northern Eurasian semi-domestic reindeer (*Rangifer tarandus*)": Supplementary_Figure.docx

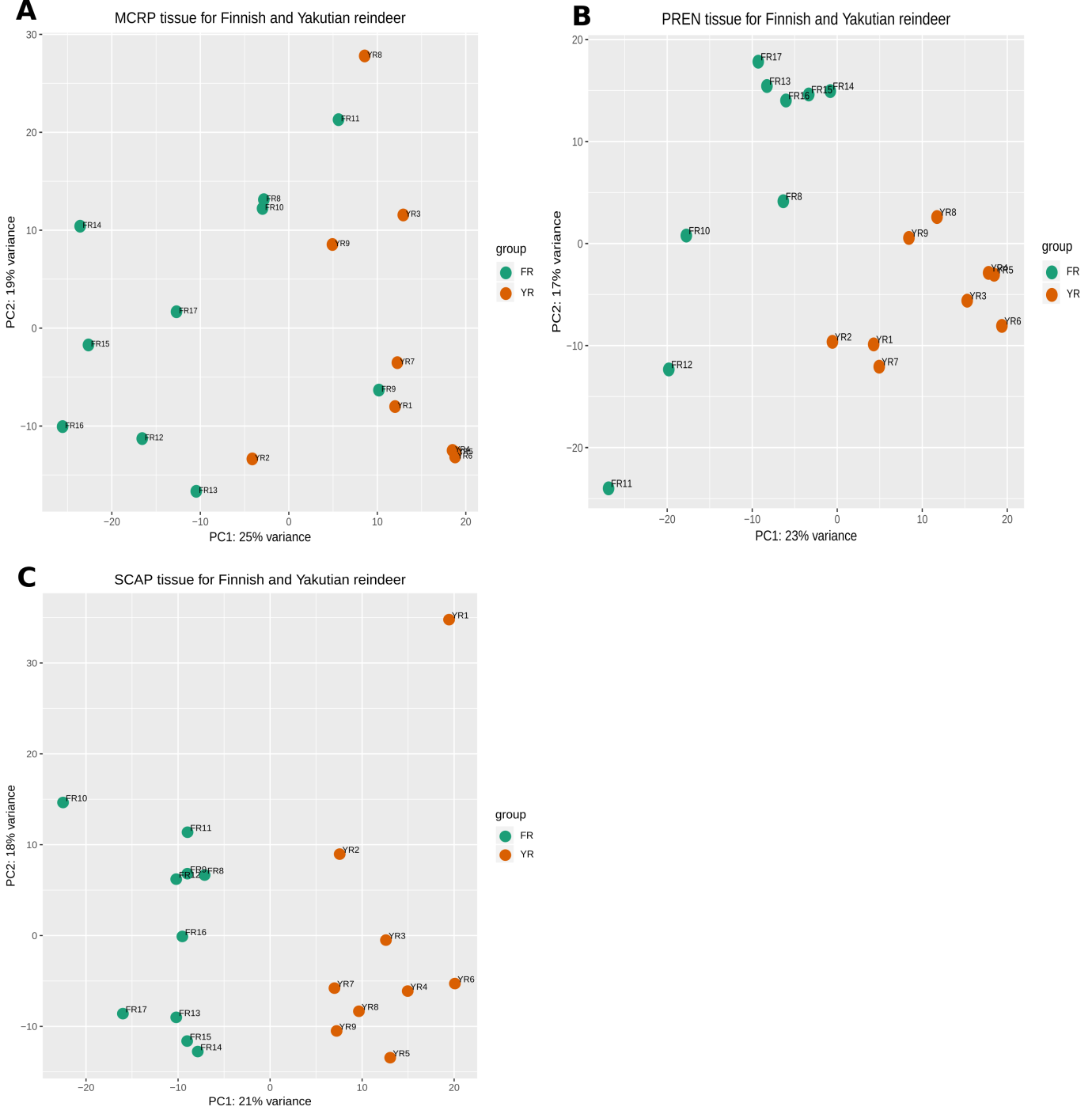

**Figure S1.** PCA plot based on region for each tissue. **(A)** metacarpal adipose tissue, **(B)** perirenal adipose tissue and **(C)** prescapular adipose tissue

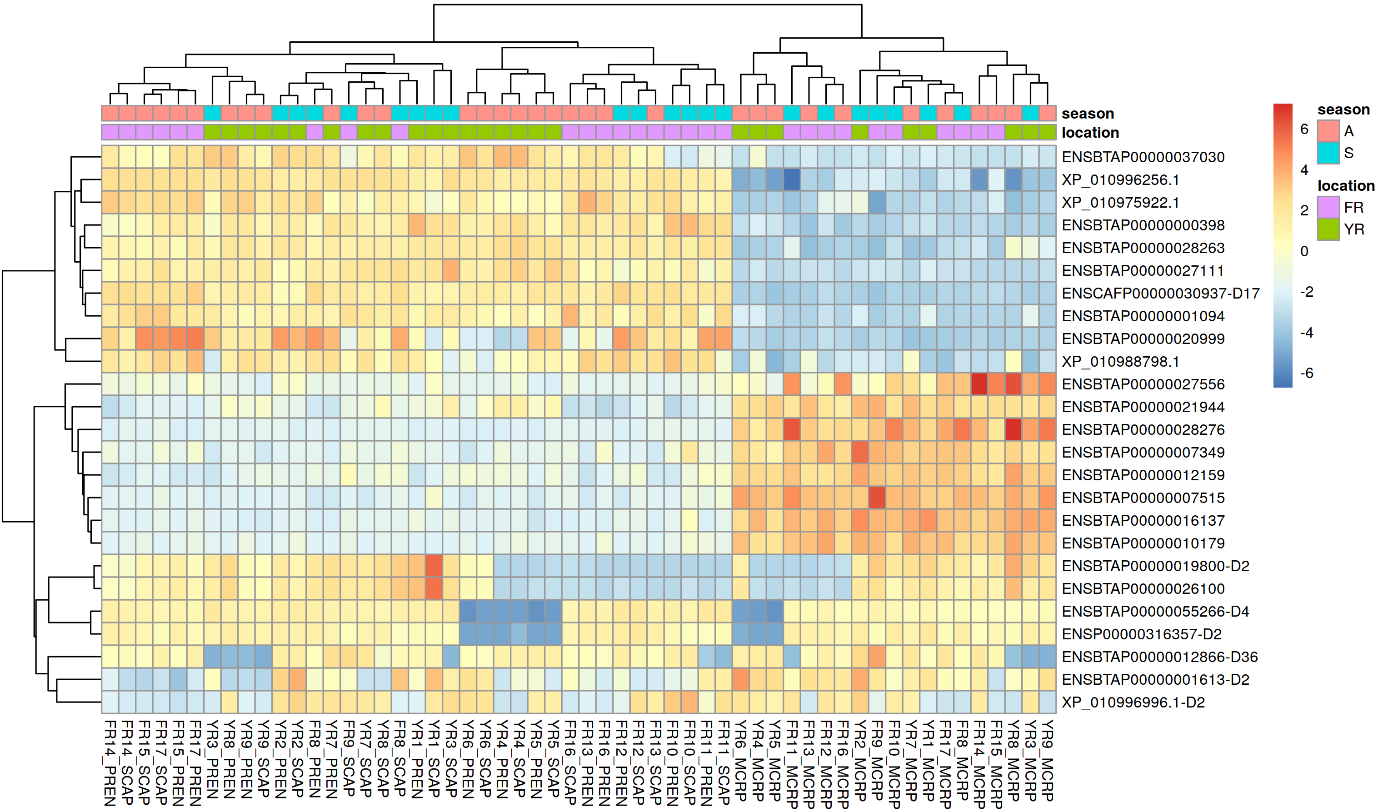

**Figure S2**. Top 25 most highly variable genes (the top 25 genes with the highest variance across samples). In the heatmap plot S stands for early spring; early winter (A); Finnish reindeer (FR) and Yakutian reindeer (YR); metacarpal tissue (MCRP); perirenal (PREN) and prescapular (SCAP).

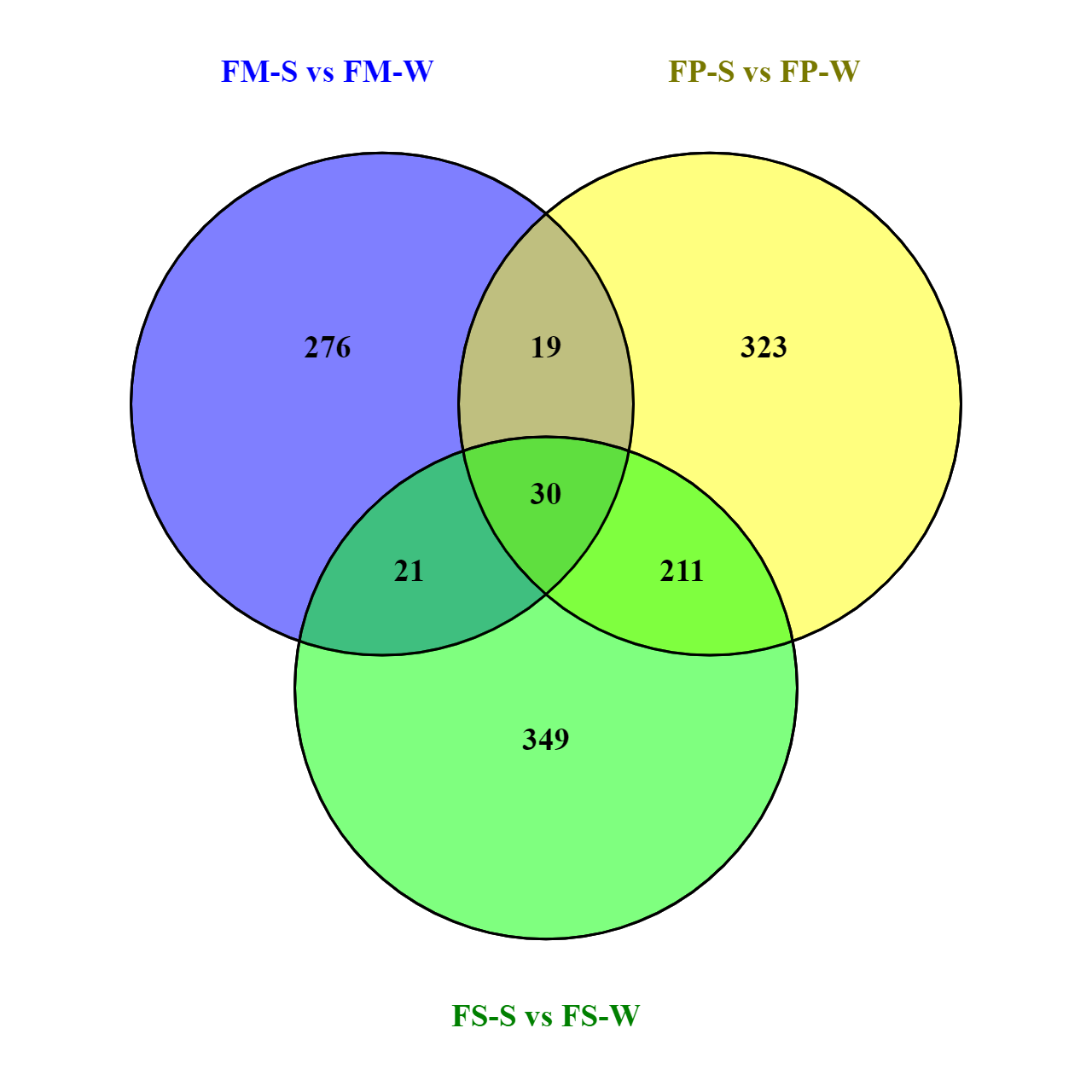

Figure S3. The number of shared and unique significant DEGs between three adipose tissues in Finnish reindeer due to seasonal differences. Significant DEGs detected in Finnish reindeer for three adipose tissues due to seasonal change: FM-S vs. FM-W, FP-S vs. FP-W and FS-S vs. FS-W.

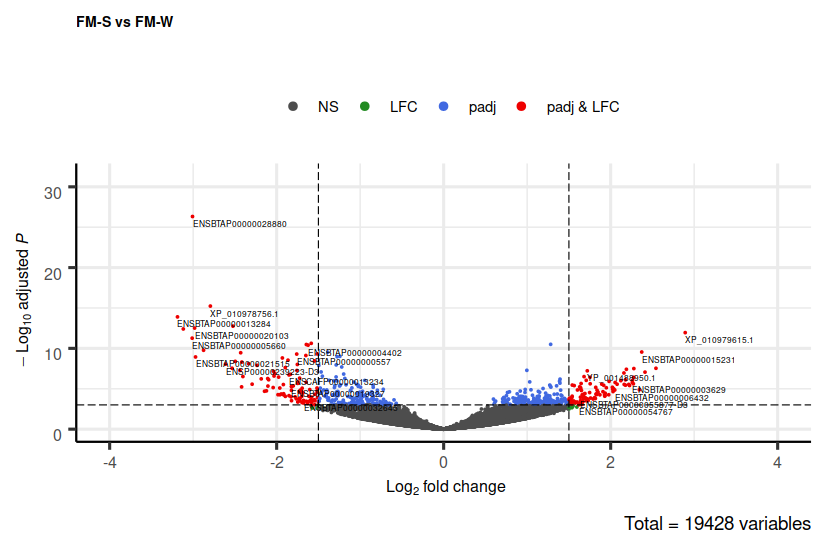

Figure S4. Volcano plot of differentially expressed genes between early spring and early winter for metacarpal adipose tissue in Finnish reindeer (FM-S vs. FM-W). The x-axis represents log2 fold change, while the y-axis represents adjusted p values (−log10). The red dots indicate the significantly differentially expressed genes (upregulated or downregulated). The threshold padj ≤ 0.05 and log2 |fold change| > 1.5 were used to identify the significant DEGs. The green dots indicate log2 |fold change| > 1.5. The blue dots indicate padj ≤ 0.05. The black dots indicate genes that did not show significant difference due to seasonal differences.

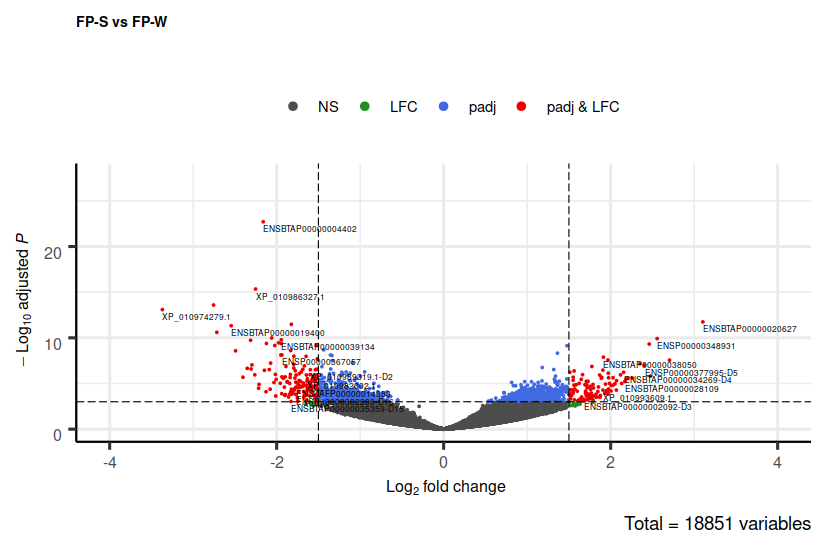

Figure S5. Volcano plot of differentially expressed genes between early spring and early winter for perirenal adipose tissue in Finnish reindeer (FP-S vs. FP-W).

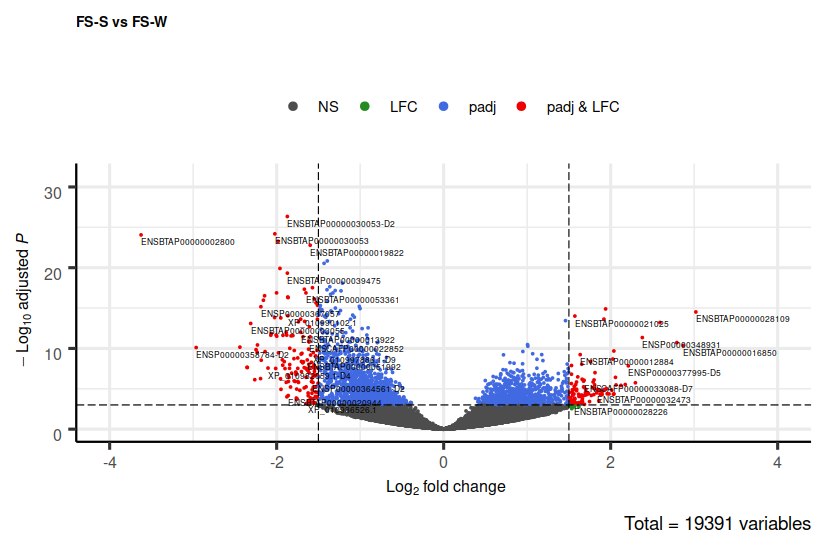

Figure S6. Volcano plot of differentially expressed genes between early spring and early winter for prescapular adipose tissue in Finnish reindeer (FS-S vs. FS-W).

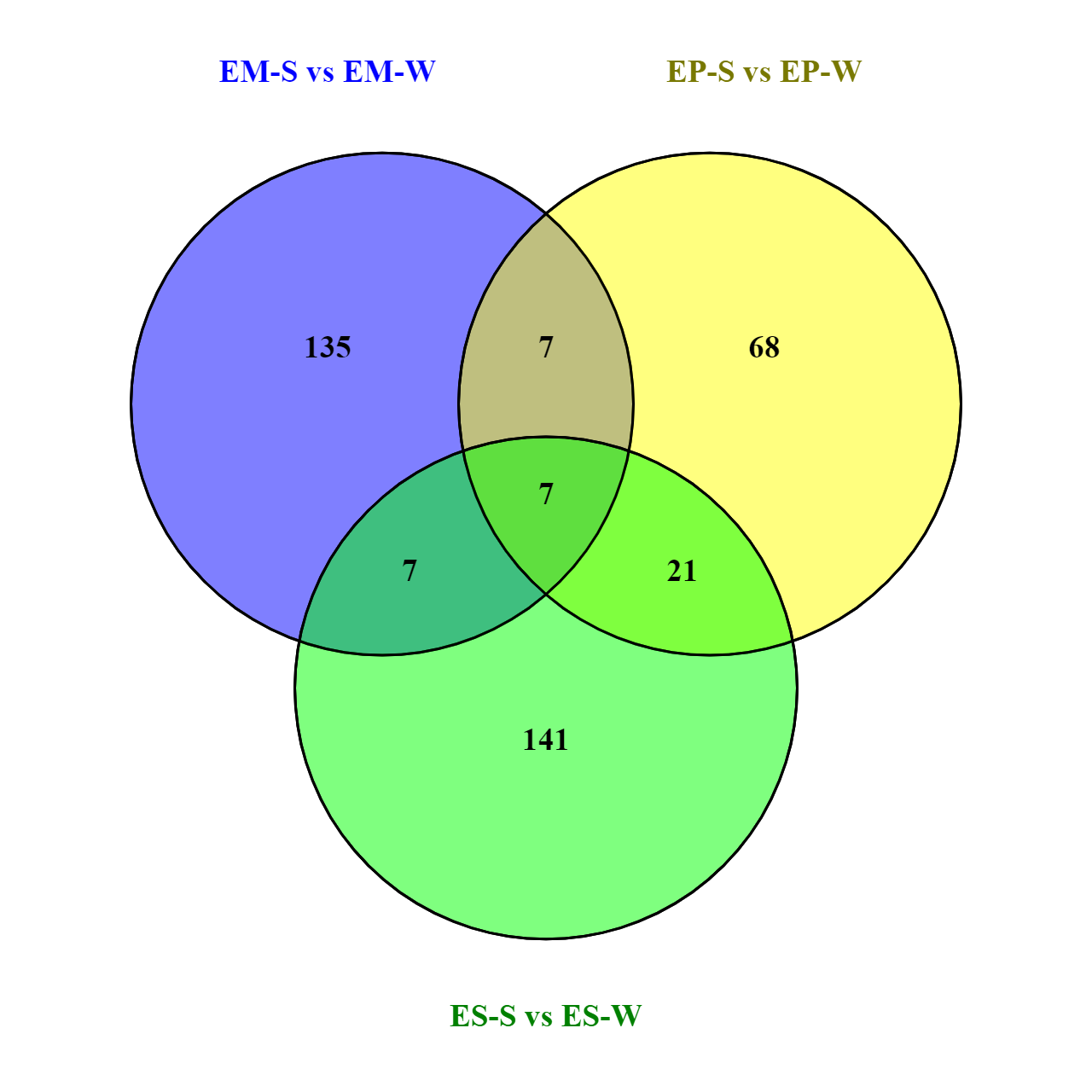

Figure S7. The number of shared and unique significant DEGs between three adipose tissues in Even reindeer due to seasonal differences. Significant DEGs detected in Even reindeer for three adipose tissues due to seasonal change (EM-S vs. EM-W, EP-S vs. EP-W and ES-S vs. ES-W).

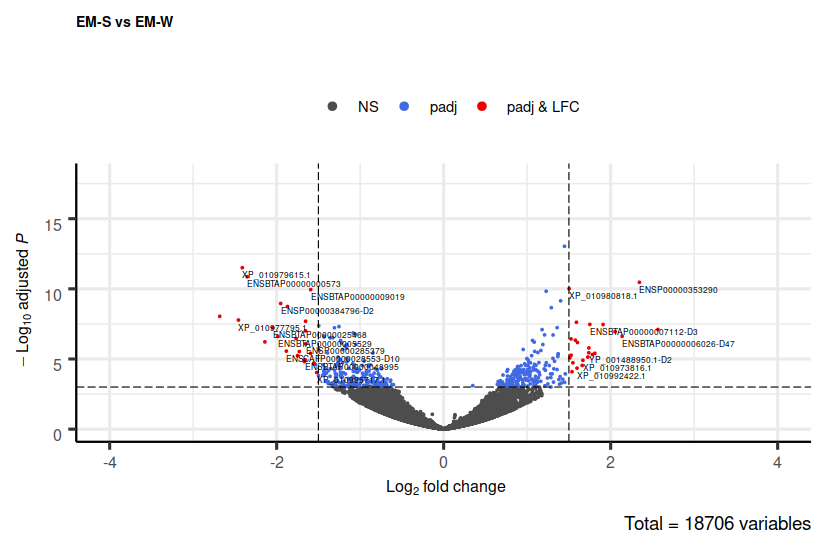

Figure S8. Volcano plot of differentially expressed genes between early spring and early winter for metacarpal adipose tissue in Even reindeer (EM-S vs. EM-W).

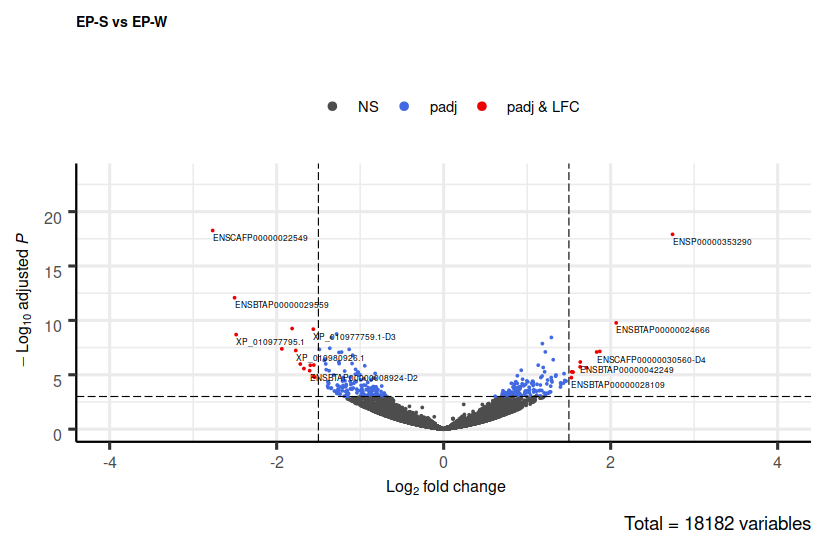

Figure S9. Volcano plot of differentially expressed genes between early spring and early winter for metacarpal adipose tissue in Even reindeer (EM-S vs. EM-W).

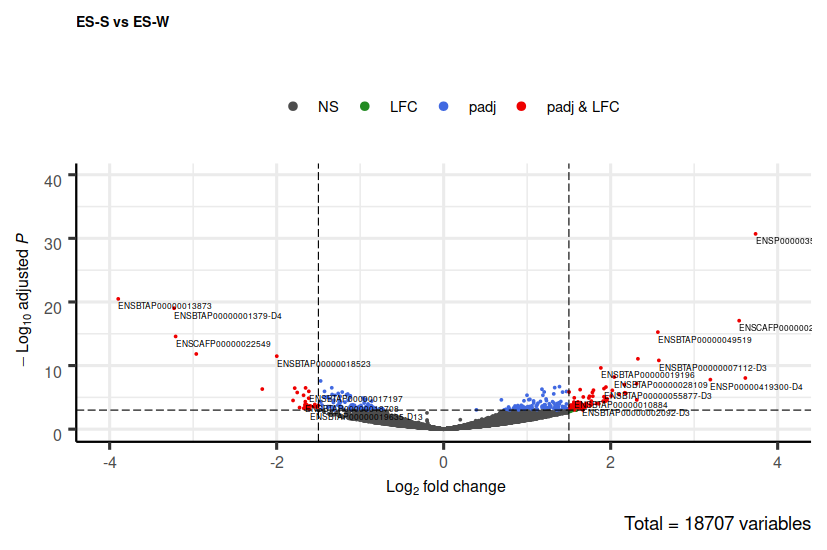

Figure S10. Volcano plot of differentially expressed genes between early spring and early winter for metacarpal adipose tissue in Even reindeer (EM-S vs. EM-W).

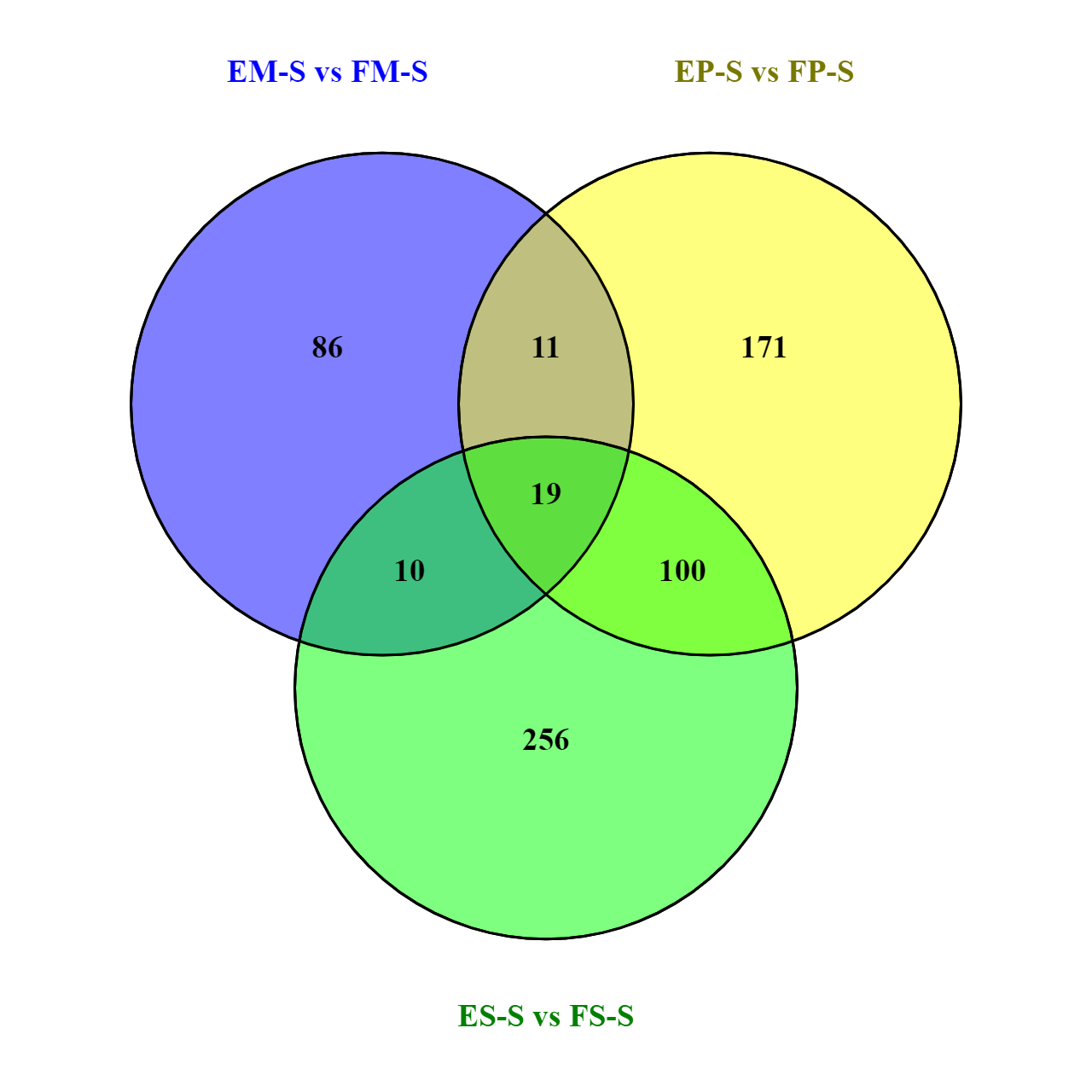

Figure S11. The number of shared and unique significant DEGs in the three adipose tissues detected between Even reindeer and Finnish reindeer due to regional difference in early spring. Significant DEGs detected in three adipose tissues due to regional difference in early spring: EM-S vs. EM-S, EP-S vs. EP-S and ES-S vs. ES-S.

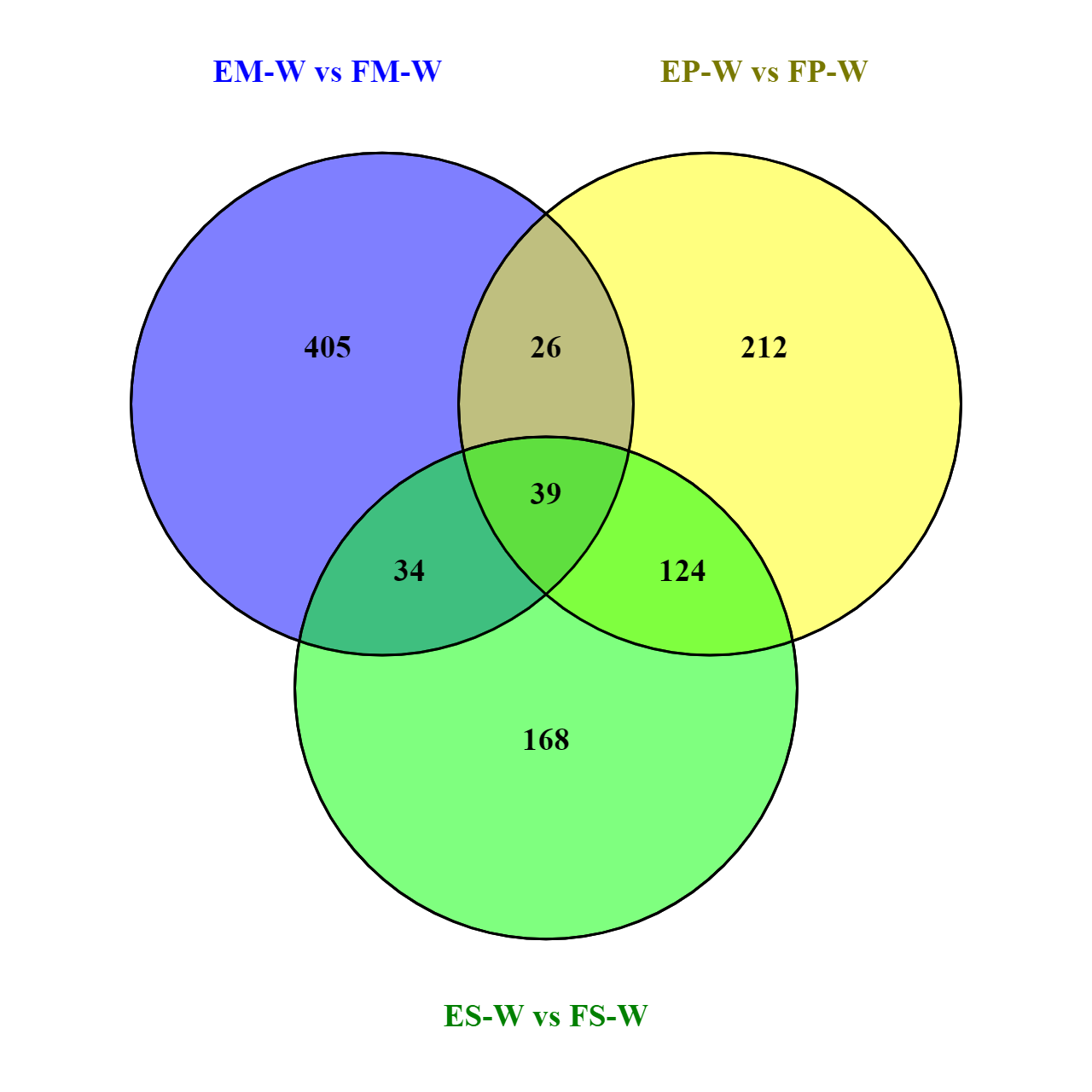

Figure S12. The number of shared and unique significant DEGs in the three adipose tissues detected between Even reindeer and Finnish reindeer due to regional difference in early winter. Significant DEGs detected in three adipose tissues due to regional difference in early winter: EM-W vs. EM-W, EP-W vs. EP-W and ES-W vs. ES-W.
